# Integrative single-cell meta-analysis reveals disease-relevant vascular cell states and markers in human atherosclerosis

**DOI:** 10.1101/2022.10.24.513520

**Authors:** Jose Verdezoto Mosquera, Gaëlle Auguste, Doris Wong, Adam W. Turner, Chani J. Hodonsky, Christian L. Lino Cardenas, Konstantinos Theofilatos, Maxime Bos, Maryam Kavousi, Patricia A. Peyser, Manuel Mayr, Jason C. Kovacic, Johan L. M. Björkegren, Rajeev Malhotra, Sander W. van der Laan, Chongzhi Zang, Nathan C. Sheffield, Clint L. Miller

**Affiliations:** Department of Biochemistry and Molecular Genetics, University of Virginia, Charlottesville, Virginia, USA.; Center for Public Health Genomics, University of Virginia, Charlottesville, Virginia, USA.; Cardiovascular Research Center, Cardiology Division, Department of Medicine, Massachusetts General Hospital, Harvard Medical School, Boston, Massachusetts, USA.; King’s British Heart Foundation Centre, King’s College London, United Kingdom.; Department of Epidemiology, Erasmus University Medical Center, The Netherlands.; Department of Epidemiology, University of Michigan, Ann Arbor, Michigan, USA.; Cardiovascular Research Institute, Icahn School of Medicine at Mount Sinai, New York, New York, USA.; Victor Chang Cardiac Research Institute, Darlinghurst, New South Wales, Australia.; St. Vincent’s Clinical School, University of New South Wales, Sydney, New South Wales, Australia.; Department of Genetics and Genomic Sciences, Icahn Institute for Genomics and Multiscale Biology, Icahn School of Medicine at Mount Sinai, New York, New York, USA.; Department of Medicine, Karolinska Institutet, Huddinge, Sweden.; Central Diagnostics Laboratory, Division Laboratories, Pharmacy, and Biomedical Genetics, University Medical Center Utrecht, Utrecht University, Utrecht, The Netherlands.; Department of Public Health Sciences, University of Virginia, Charlottesville, Virginia, USA.

## Abstract

Coronary artery disease (CAD) and atherosclerosis are characterized by plaque formation in the arteries wall. CAD progression involves complex interactions and phenotypic plasticity within and between distinct vascular and immune cell lineages. Single-cell RNA-seq (scRNA-seq) studies have highlighted lineage-specific transcriptomic signatures, but the reported cell phenotypes in humans remain controversial. Here, we meta-analyzed four scRNA-seq datasets, creating the first map of human cell diversity in atherosclerosis. We generated an atlas of 118,578 high-quality cells, characterized cell-type diversity and provided insights into smooth muscle cell (SMC) phenotypic modulation, transcription factor activity and cell-cell communication. We integrated genome-wide association study (GWAS) data and uncovered a critical role for modulated SMC phenotypes in CAD and coronary calcification. Finally, we identified candidate markers of fibromyocyte and fibrochondrogenic human SMCs (*LTBP1* and *CRTAC1*) that may serve as proxies of atherosclerosis progression. Altogether, we created a unified cellular map of atherosclerosis informing cell state-specific mechanistic and translational studies of cardiovascular diseases.

## Introduction

Cardiovascular diseases, such as coronary artery disease (CAD), are the leading global causes of mortality and morbidity^1^. The pathological hallmark of CAD is atherosclerosis, a chronic build- up of plaque inside arterial walls, which can lead to thrombus formation and myocardial infarction (MI) or stroke^2–5^. This process involves a complex interplay of both immune and vascular cell types and cell state transitions along a continuum^6, 7^. In response to injury of the inner vessel wall layer, contractile smooth muscle cells (SMCs) transition to a more proliferative and migratory state^8, 9^ and endothelial cells to a mesenchymal state in early and advanced atherosclerosis^10, 11^. Thus, a thorough assessment of cell heterogeneity and plasticity within the vessel wall is paramount to uncover new knowledge regarding atherosclerosis development and progression.

The advent of single-cell sequencing technologies has enabled study of gene expression and regulation in disease and development at the single-cell level. For instance, single-cell RNA sequencing (scRNA-seq) studies have resolved the cellular diversity and gene signatures in human and murine atherosclerotic lesions^12–16^ as well as in non-lesion arteries^17^. By combining lineage tracing and scRNA-seq, studies have shown that SMCs readily transform into a multipotent “pioneer” cell type in response to pro-atherogenic stimuli^18–20^. However, their fate after this transition remains controversial, with a few studies agreeing they can become fibroblast-like (fibromyocytes)^18^ or osteogenic-like (fibrochondrocytes; FCs)^19^, while others suggest they adopt pro-inflammatory or macrophage-like properties^8, 20^. Biological interpretation of these individual studies could be potentially confounded by limited sample sizes, experimental design, or other technical factors. Thus, there remains a need for a consensus single-cell reference^21–23^ that spans atherosclerotic disease stages in humans.

Here, we harmonize and meta-analyze four single-cell studies of human atherosclerosis, encompassing both early and advanced lesion and non-lesion samples (**Fig. 1a and Supplementary Table 1**). This high-resolution atlas of 118,578 high quality cells enables the discovery of previously missed vascular and immune cell types and clarifies markers for known disease-relevant immune cells (e.g., inflammatory, and foamy macrophages). We perform integrative downstream analyses and GWAS trait enrichment to define cardiovascular traits and disease-relevant etiologic cell types and states. We further validate SMC phenotypes identified in lineage-tracing studies, reveal underrepresented SMC states from individual scRNA-seq studies and highlight *CRTAC1 and LTBP1* as new candidate markers of synthetic and pro-calcifying SMCs and plaque stability in humans. This comprehensive map of cell diversity in human atherosclerosis provides a critical step towards translating mechanistic knowledge and developing more targeted interventions.

**Figure 1.**
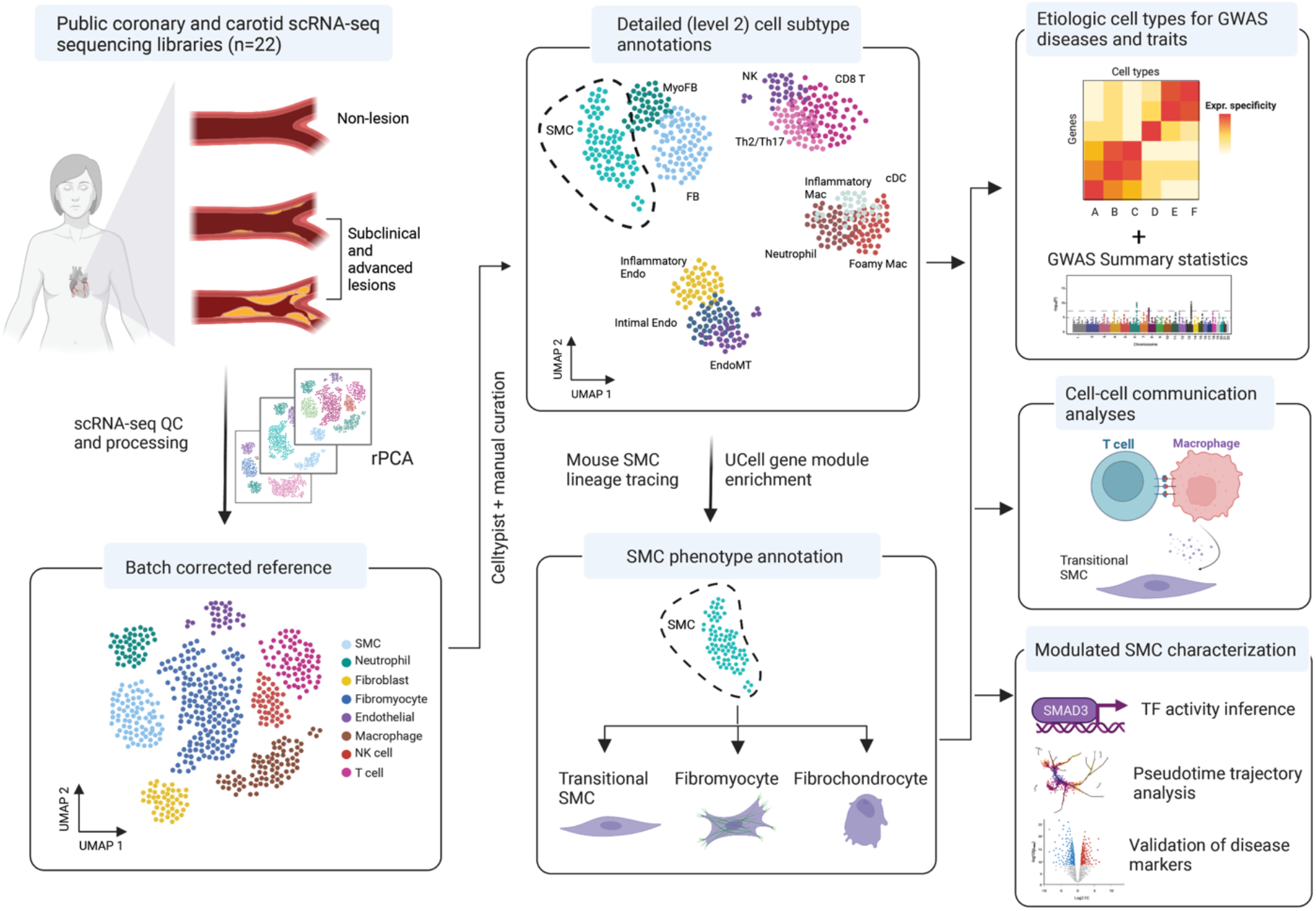
General workflow of the study. We collected human atherosclerosis scRNA-seq libraries from four publications, including three from atherosclerotic lesions of varying stages^16, 18, 19^ and one with no CAD diagnosis or discernable lesions^17^. After uniform QC and processing of each library, we benchmarked four batch-correction methods. We then used transfer learning as well as machine learning classifiers and literature markers to define broad (level 1) and granular (level 2) cell type annotations across vascular and immune lineages. We also leveraged murine lineage-traced smooth muscle cell (SMC) gene modules to identify modulated SMC populations in human data. We then analyzed the resulting integrated scRNA map in several ways: integration of GWAS data for identification of etiologic cell types, cell communication analyses, pseudotime inference, TF activity predictions and identification of candidate novel human-specific gene markers. See **Methods** for details.

## Results

### Integration of lesion and non-lesion artery datasets

We sought to build a comprehensive well-powered single-cell reference to further investigate complex vascular processes such as SMC phenotypic modulation. To achieve proper representation of mural cells and immune types and to span the continuum of CAD risk, we used four human datasets including atherosclerotic lesions (*Wirka* et al.^18^, *Pan* et al.^19^, and *Alsaigh* et al.^16^), and from non-lesion coronary arteries^17^ (**Supplementary Table 1**).

We established a standardized pipeline for quality control (QC; **Supplementary Fig. 1** see Methods^24–27)^ and uniform processing of the 22 raw sequencing libraries, and observed improved separation and cohesion of cell clusters post-filtering (**Supplementary Fig. 2**). To find the optimal integration approach, we benchmarked top-performing integration tools **(Methods)**^26, 28–31^ on a subset of the libraries^16, 18, 19^ and found that reciprocal PCA (rPCA) and Harmony outperformed the other tools in terms of running time. We also evaluated the effectiveness of batch removal, conservation of biological variation and clustering purity **(Methods)**^32^. We found that rPCA achieved the best balance in terms of running time, batch mixing, separation of unique cell types and clustering purity across the tested granularity parameters. Integration of libraries with rPCA yielded a total of 118,578 high-quality cells and 41 Louvain clusters. To further ensure proper batch removal, we inspected UMAP embeddings of the reference and found mixing across the four studies for the major cell compartments (**Supplementary Fig. 2**).

We next annotated our clusters using a two-level system. We labeled cells in UMAP space with broad terms or major lineage names (level 1) and then provided a more granular resolution of annotations combining manual and automated approaches (level 2; Methods). We defined level 1 annotations by reprocessing and transferring cell type labels from the Tabula Sapiens (TS) vasculature single-cell atlas^33^. We found that labels were assigned with remarkably high confidence scores (**Supplementary Fig. 3**). These annotations were supported by the expression of well-established marker genes in corresponding level 1 clusters (**Fig. 2a-b**) and confirmed that batch effects had been properly removed while conserving biological variation.

**Figure 2.**
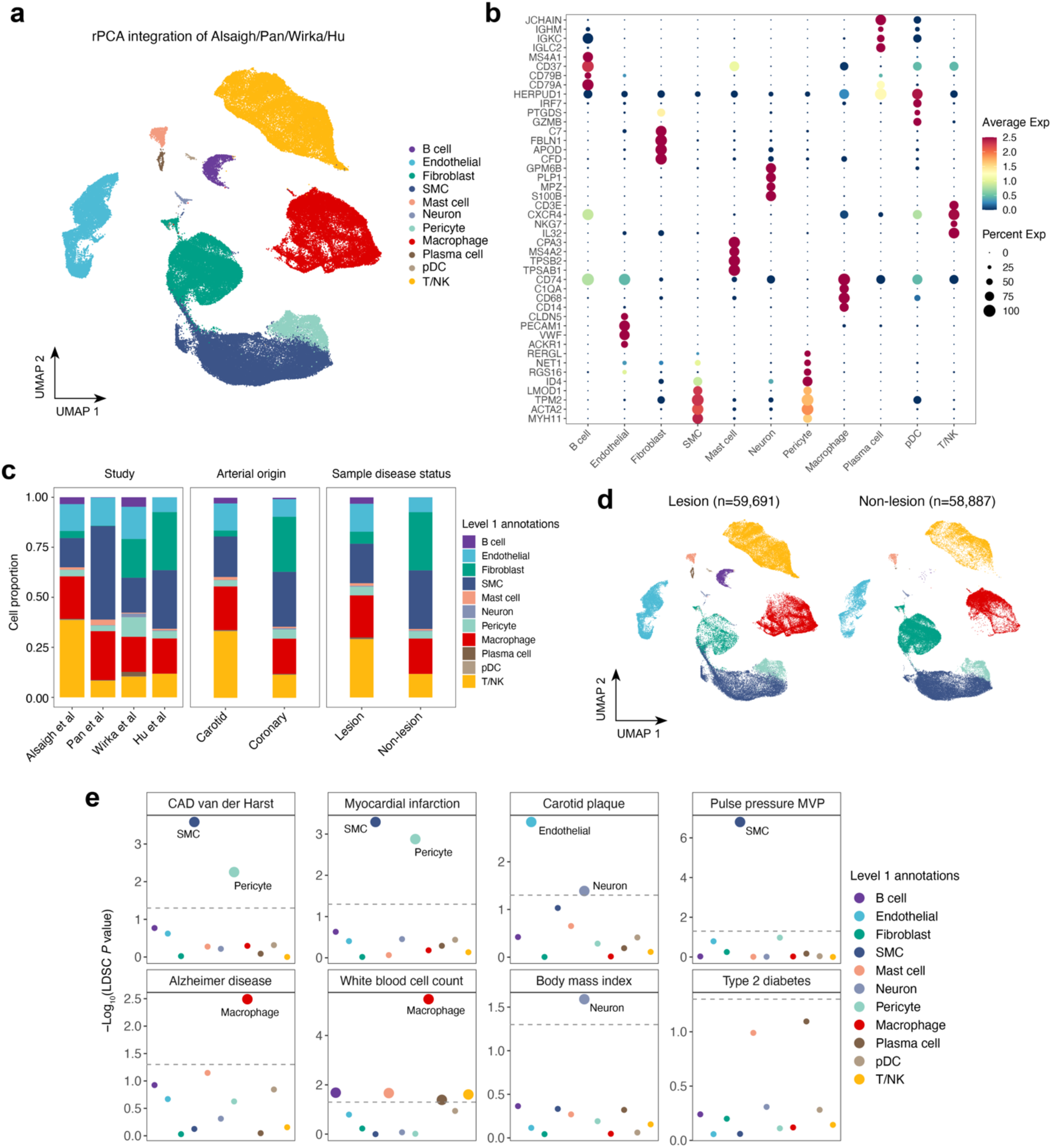
Integration of single cell data identifies major cell compartments in atherosclerosis. (**a**) UMAP representation of 118,578 cells based on rPCA integration of 22 sequencing libraries. Dot colors depict broad cell lineage labels (level 1). (**b**) Dot plot of top five marker genes SCTransform-normalized expression by major cell lineage compartment. Dot size represents the portion of cells expressing the gene per level 1 compartment. (**c**) Stacked bar plot showing the distribution of level 1-annotated cells across included publications, arterial beds (coronary, carotids), and lesion status (lesion, non-lesion). (**d**) UMAP representation of level1-annotated cells across lesion status. (**e**) LD Score Regression applied to gene expression specificity (LDSC-SEG) analyses prioritizing the contribution of level 1-annotated cell type to cardiovascular and non-cardiovascular GWAS traits. LDSC analysis was carried out using a gene expression specificity matrix generated with CELLEX^103^. Large circles depict cells that passed the cutoff of FDR < 5% at -log10(*P*) = 1.301.

We observed a balanced number of cells labeled as macrophages and endothelial cells across studies. However, there were slightly more SMCs in *Pan et al* and T/NK cells in *Alsaigh et al* and slight biases from small clusters (e.g., plasma cells, B cells) between studies (**Fig. 2c**). This shows that individual studies may under-represent key cell types. We also observed overrepresented fibroblasts from coronary datasets (*Wirka* et al. and *Hu* et al.), as expected given the intact coronary vessel wall layers compared to carotid plaques. When comparing cell type frequencies across disease status, we observed a greater proportion of B cells, plasma cells and pDCs in lesion samples **(Fig. 2c-d).** Further, libraries from *Alsaigh et al* had the highest proportion of T cells among all studies, consistent with the advanced stage of the carotid lesions (**Supplementary Fig. 3**).

Differentially expressed genes for each cluster **(Methods and Supplementary Table 2)** were used to run gene ontology (GO) enrichment analyses. We found enrichment for terms in their expected clusters, such as “muscle contraction” in SMCs and “endothelial development” in endothelial cells (ECs). We also observed over-representation of terms such as “extracellular matrix organization” in SMCs, likely due to the presence of phenotypically modulated SMCs^9^ (**Supplementary Fig. 3, Supplementary Table 3**). In contrast, myeloid and lymphoid clusters were enriched for immune-related terms such as “antigen processing and presentation” and “regulation of T cell activation”. These results confirmed the accuracy of our level 1 annotations.

### Defining candidate etiologic cell types for complex traits

Next, we identified etiologic cell types enriched for atherosclerosis-related traits using our level 1 cell type annotations by performing LD score regression applied to specifically expressed genes (LDSC-SEG) analyses^34^ using GWAS summary statistics for cardiovascular disease (CVD) and non-CVD traits as described^35–40^ (**Methods**). SMC and pericyte gene signatures were significantly enriched (FDR < 0.05) for CV traits such as pulse pressure, CAD, and MI (**Fig. 2e and Supplementary Table 4**). EC signatures were enriched for carotid plaque associations (**Fig. 2e**). Consistent with previous studies^34, 41^, we observed macrophages were highly enriched for Alzheimer’s disease and white blood cell count GWAS signals. We also found high enrichment of neurons for body mass index (BMI). These findings highlight the value of integrating single-cell and human genetic data to discover atherosclerosis trait-relevant cell types, such as SMC and ECs.

### Defining cell subtype heterogeneity in human atherosclerosis

Next, we surveyed the 41 clusters using a combination of automated and manual annotation **(Methods)**. Manual annotations included markers of lymphoid, myeloid and endothelial cell subtypes from the literature^21–23, 42–47^. We then verified manual annotations using the CellTypist machine learning classifier^48^ resulting in a more granular map of cell diversity in human atherosclerosis **(Fig. 3a).** We summarize some of the most representative cell subpopulations and markers **(Supplementary Table 5)** below:

**Figure 3.**
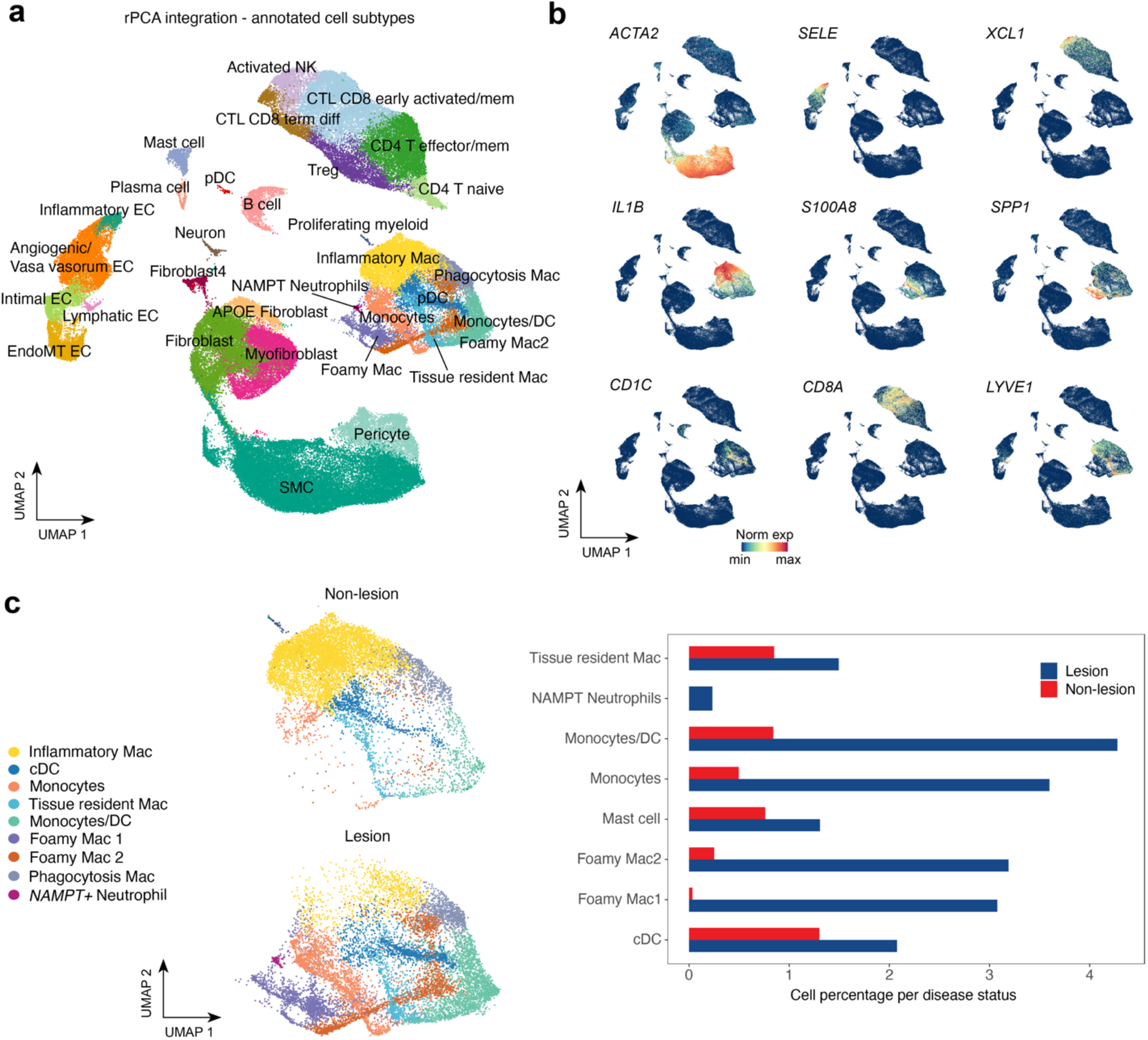
Human atherosclerosis cell subpopulations (level 2) and distribution of myeloid subtypes across disease status. (**a**) UMAP representation of cell subtypes (level 2 labels) within the largest level 1 cell compartments (T/NK, Macrophages, Endothelial, Fibroblast). (**b**) UMAP plots of top marker/canonical marker genes delineating level 1 annotations. SCTransform-normalized gene expression is indicated by color. (**c**) UMAP and bar plot of level 2 Myeloid cell subtypes according to lesion status. Frequencies for each subtype shown in the bar plot are normalized to the total number of cells in each condition (lesion n=59691; non-lesion n=58887) and shown as percentages.

*Endothelial diversity:* We identified cells highly expressing classical endothelial markers (*PECAM1*, *CLDN5)* relative to neighboring clusters (**Fig. 3a**). Intimal ECs were defined by expression of homeostatic EC marker genes (e.g., *RAMP2)*^49^. We also identified a cluster of pro-angiogenic ECs with upregulated vasa vasorum or angiogenic genes (e.g., *ACKR1, AQP1 and FABP4)*^50–52^. Adjacent, we identified a cluster with elevated expression of chemokine and adhesion molecules (*SELE*, *CCL2*) (**Fig. 3a-b**), likely reflecting a pro-inflammatory state^53^. EndoMT ECs^54^ were defined by the expression of ECM genes (*COL1A2, FN1)* and contractile genes. Finally, we defined a small subcluster of lymphatic ECs based on the expression of *LYVE1* and *CCL21*^55^.

*Myeloid diversity:* We identified a subset of myeloid cells, inflammatory macrophages expressing known “M1-polarized” macrophage markers (e.g., *IL1B*, *TNF*) (**Fig. 3a-b**). Foamy macrophages were annotated by higher expression of lipid genes (*APOE* and *FABP5)* and lower inflammatory genes^15, 21, 23^. We also identified resident macrophages (*LYVE1, FOLR2*), classical monocytes (*S100A8*, *S100A9, LYZ*), and conventional dendritic cells (*CD1C*, *CLEC10A*)^56–58^. Importantly, we resolved critical smaller myeloid populations including plasmacytoid dendritic cells (pDCs)^59, 60^ and neutrophils (*NAMPT*, *S100A8*), overlooked by previous individual human scRNA-seq datasets. Consistently, we found more prevalent monocytes, foamy macrophages, and other myeloid populations in lesion-containing libraries (**Fig. 3c**).

*Lymphoid diversity:* We identified Natural Killer (NK) and several subpopulations of T cells based on differential expression of *CD69*^61^, *CD8A/B*, among others **(Fig 3a-b, Supplementary Fig. 3)**. We identified markers of early activated and memory/naive CD8 cytotoxic T cells (CTLs)^62, 63^, terminally differentiated CTLs, Th17 and Th2 helper cells and regulatory T cells **(Supplementary Fig. 3)**. Finally, we defined clusters of B cells (*CD79A*, *CD79B*) and plasma cells (*IGLC2*, *IGHM*, *JCHAIN*). While all lymphoid populations showed larger frequencies in lesions, we found that B cells, plasma cells and pDCs were highly depleted in non-lesion libraries (**Supplementary Fig. 3**).

*Fibroblast diversity:* Defining fibroblast diversity in atherosclerosis is particularly challenging given the low specificity of widely used fibroblast markers^46^. We found that most cells in this compartment express traditional fibroblast ECM markers such as *LUM* and *DCN* (**Supplemental Table 5**). We were able to dissect a subset of fibroblasts that upregulated the contractile marker *ACTA2* (**Fig. 3b**) as well as complement genes (*C3* and *C7*). This subset likely represents activated fibroblasts (myofibroblasts) known to adopt increased contractile, ECM-producing, and pro-inflammatory states in response to injury or atherosclerotic stimuli^46, 54^. We also identified a group of cells within the fibroblast compartment strongly expressing *APOE* in addition to the chemokine ligands *CXCL12* and *CXCL14* and complement genes, which we term APOE fibroblasts.

### Characterization of SMC phenotypes in human atherosclerosis

To refine the role of human SMC phenotypes, we next compared our human scRNA reference to a recent scRNA meta-analysis of murine vascular SMCs^22^. First, we subsetted the full human atlas to include only SMCs, pericytes and a subset of fibroblasts. We then assessed enrichment of lineage-traced murine SMC gene modules on a per-cell basis using the UCell R package^64^(**Methods**). This showed a progressive loss of the murine SMC contractile signature within a portion of the human subset, coincident with a gain in the *Lgals3*+ transitional gene signature (**Fig. 4a)**, supporting a transitional SMC signature in humans. Further, we detected an enriched signature of the murine calcification-promoting fibrochondrocytes distinct from non-SMC-derived fibroblasts. DE markers for each cluster and UCell module enrichment scores were used as a guide to annotate SMCs as contractile, ECM-rich transitional SMCs, fibromyocytes and fibrochondrocytes (FCs) (**Fig. 4b-c, Supplementary Fig. 4, Supplementary Table 6**).

**Figure 4.**
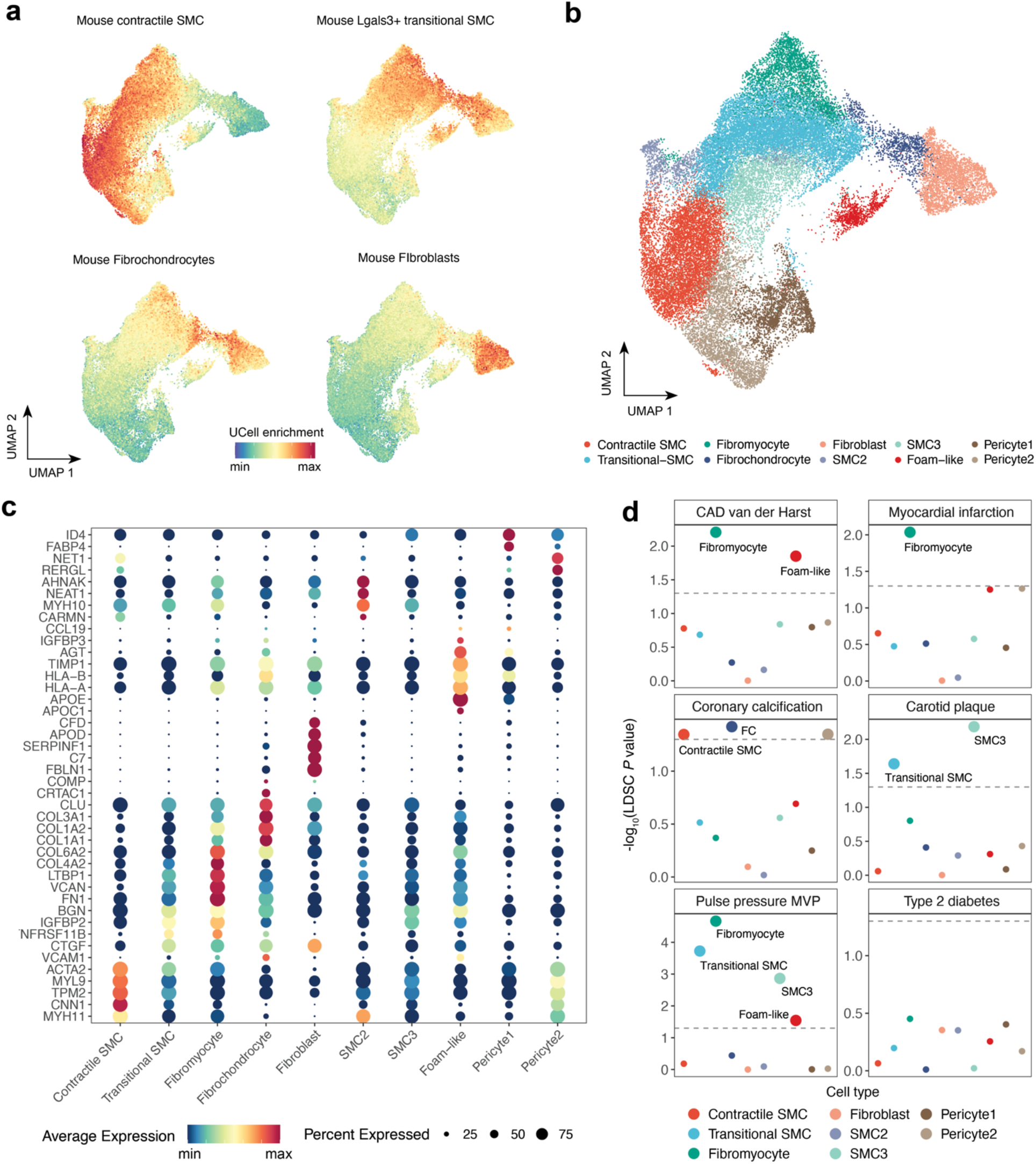
Characterization of etiologic SMC phenotypes for cardiovascular traits and diseases. (**a**) UCell enrichment of meta-analyzed SMC murine gene modules (Contractile, *Lgals3+* transitional, Fibrochondrocytes) and non-SMC-derived fibroblasts in the level 1 SMC compartment as well as a Fibroblast subset. UCell^64^ scores were calculated for each cell based on the Mann-Whitney U statistic where higher scores depict a higher enrichment for the tested gene signature. (**b**) UMAP embeddings of SMC level 2 labels (described in **a**) in addition to pericytes and a subset of Fibroblasts. Labels were defined using UCell scores as reference for SMC differentiation state in addition to DE markers from Louvain clusters at a resolution=0.9. (**c**) Dot plot showing expression of top marker genes after SCTransform-normalization for SMC level 2 labels. Dot size represents the portion of cells expressing the gene. (**d**) LD score regression applied to specifically expressed genes (LDSC-SEG) analyses prioritizing the contribution of SMC phenotypes, Pericytes, and Fibroblasts to cardiovascular GWAS traits. Type 2 diabetes was used as a negative control in this analysis. LDSC was carried out using a gene expression specificity matrix for SMC clusters generated with CELLEX^103^. Large circles depict cells that passed the cutoff of FDR < 5% at -log10(*P*) = 1.301.

We observed similar proportions of contractile, transitional SMCs and fibromyocytes across arterial beds and lesion status, consistent with previous reports^65^. However, in line with their role in calcification, FCs predominated in lesions compared to non-lesion samples. The FC annotation was further supported by higher ESμ values for SOX9 and RUNX2 **(Supplementary Fig. 4)**, known master regulators of SMC osteochondrogenic transitions^66^. At a global level, SMCs, transitional SMCs, fibromyocytes, and FCs were enriched for relevant biological processes thus validating our annotation approach (**Supplementary Table 7**). Interestingly, we also identified a cluster enriched for the transitional SMC gene signature and expressing lipid metabolism genes (*APOE*, *APOC1, AGT)*, which we termed “foam-like” SMCs (**Fig. 4b-c, Supplementary Fig. 4, Supplementary Table 6**). These cells also expressed ECM-remodeling genes such as *TIMP1* and pro-inflammatory genes *CCL19*, *CCL2, IGFBP3*, consistent with a potential role in leukocyte recruitment^67^.

Finally, we leveraged these granular SMC labels to dissect the disease relevance of SMC modulated phenotypes using LDSC-SEG. Fibromyocytes and foam-like SMCs were highly enriched for CAD heritability, while fibromyocytes were enriched for MI and subclinical CAD traits (**Fig. 4d and Supplemental Table 8**). In contrast, we observed FCs enriched for coronary artery calcification (CAC) using our recent meta-analysis summary data^68^, which further supports the robustness of this annotation. This enrichment is also consistent with our understanding of the biology of CAC, but to our knowledge has not been previously reported in any integrative single-cell and human genetic analysis.

### Cell crosstalk in human atherosclerosis

To interrogate paracrine signaling in atherosclerosis, we dissected cellular crosstalk from our level 1 and 2 annotations across lesion status using CellChat^69^. We observed strong interactions between SMCs and fibroblasts in non-lesion samples, while SMC and EC interactions with macrophages and T/NK were stronger in lesions (**Fig. 5a**). Differential signaling pathway enrichment analysis between SMCs and Macrophages revealed tumor necrosis factor alpha (TNFa) and platelet-derived growth factor (PDGF) signaling pathways enriched in non-lesion samples, whereas tumor-necrosis factor-like weak inducer of apoptosis (TWEAK) and osteopontin (SPP1) mediated signaling pathways were enriched in lesion samples (**Fig. 5b, Supplementary Table 9**). Signaling involving SPP1, specifically targeted SMCs and was mostly driven by macrophage foam cell clusters (**Fig. 5c**). This is consistent with previous studies implicating these signals in myeloid-SMC mediated vascular inflammation and calcification.

**Figure 5.**
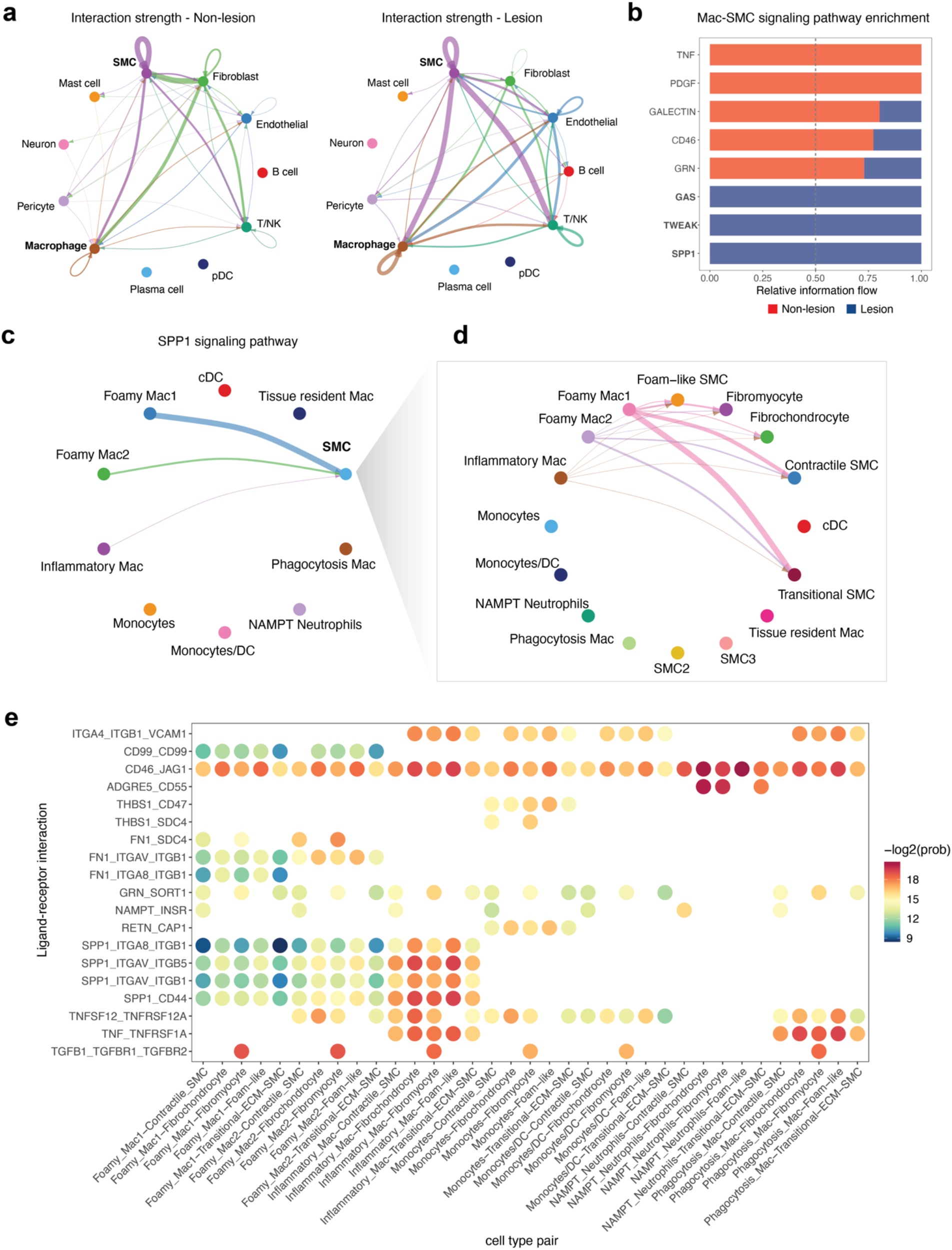
Summary of cell crosstalk in human atherosclerosis. (**a**) Circle plots depicting aggregated cell-cell communication network for level 1-labeled cell compartments leveraging the CellChat^69^ human database. Interactions considered include secreted signaling, ECM-receptor and cell-cell contacts. Interactions were calculated separately across disease status (lesion vs non-lesion). Top 30% of interactions are shown in the plot. (**b**) Stacked bar plot showing conserved and disease status-specific signaling pathways. Signaling enrichment is based on changes on pathways information flow (defined by the sum of communication probability among all pairs of cell groups in the inferred network or total weights in the network). Pathways in bold denote those that showed statistically significant (*P* < 0.05) enrichments in each disease condition. (**c**) Circle plot depicting sources and targets for SPP1 signaling using level 2 labels for myeloid cells and level 1 SMC labels. (**d**) Circle plot depicting SPP1 signaling sources and targets for level 2 Myeloid and SMC labels. (**e**) Summary dot plot of ligand-receptor interactions for level 2 myeloid and SMC annotations. Myeloid subtypes were defined as signaling sources while SMCs were defined as signaling targets. Width of the edges in the circle plot depicts the weight/strength of the interactions in (**a**,**c**,**d**).

Leveraging our SMC subtype annotations, we observed more TWEAK mediated interactions between contractile/transitional SMCs and distinct myeloid subtypes in lesion samples (**Supplementary Fig. 5**). Instead, SPP1 signals from foamy macrophage cells (foamy mac1) specifically targeted contractile and transitional SMCs (**Fig. 5d**). Using ligand-receptor interaction analyses, we also found that myeloid expressing genes encoding SPP1 ligand preferentially signal to fibromyocyte and foam-like SMC via the heterodimeric ITGA8/ITGB1 receptor (**Fig. 5e**, **Supplementary Fig. 5**, **Supplementary Table 10**).

### Modeling of SMC gene expression across pseudotime

Current evidence suggests that SMCs transition into fibromyocytes/fibrochondrocytes through an *Lgals3*+ transitional state^19, 20^. We modeled SMC de-differentiation via pseudotime analysis using Monocle 3^70^, in which we defined *MYH11*-expressing contractile SMCs as the starting point of phenotypic modulation (**Fig. 6a**). The inferred trajectory revealed a branchpoint where transitional SMCs could adopt either a fibromyocyte or FC fate. In addition, we observed more FCs in lesion samples towards the latter pseudotime stages, consistent with calcification in advanced lesions (**Fig. 6a-b**). We also identified modules specific to transitional SMC (Modules 5 and 10), fibromyocytes (Module 4) and FCs (Module 9) (**Supplementary Fig. 6, Methods**). Transitional SMC modules harbored genes involved in early SMC investment in atherosclerotic lesions (e.g., *LGALS3*)^20^, as well as cell division and proliferation (e.g., *TUBA1B* and *SIRT6*)^71^ and ECM remodeling (e.g. *KRT8* and *SPARC*). As expected, fibromyocyte module 4 included known markers (e.g., *FN1*, *VCAN*, *COL4A1/2*, *PDGFRB*). In contrast, the FC module 9 harbored chondrocyte related genes such as *BMP4*^72^, *WISP2*, and *SPRY1*^73^ in addition to known ECM genes *LUM* and *DCN*.

**Figure 6.**
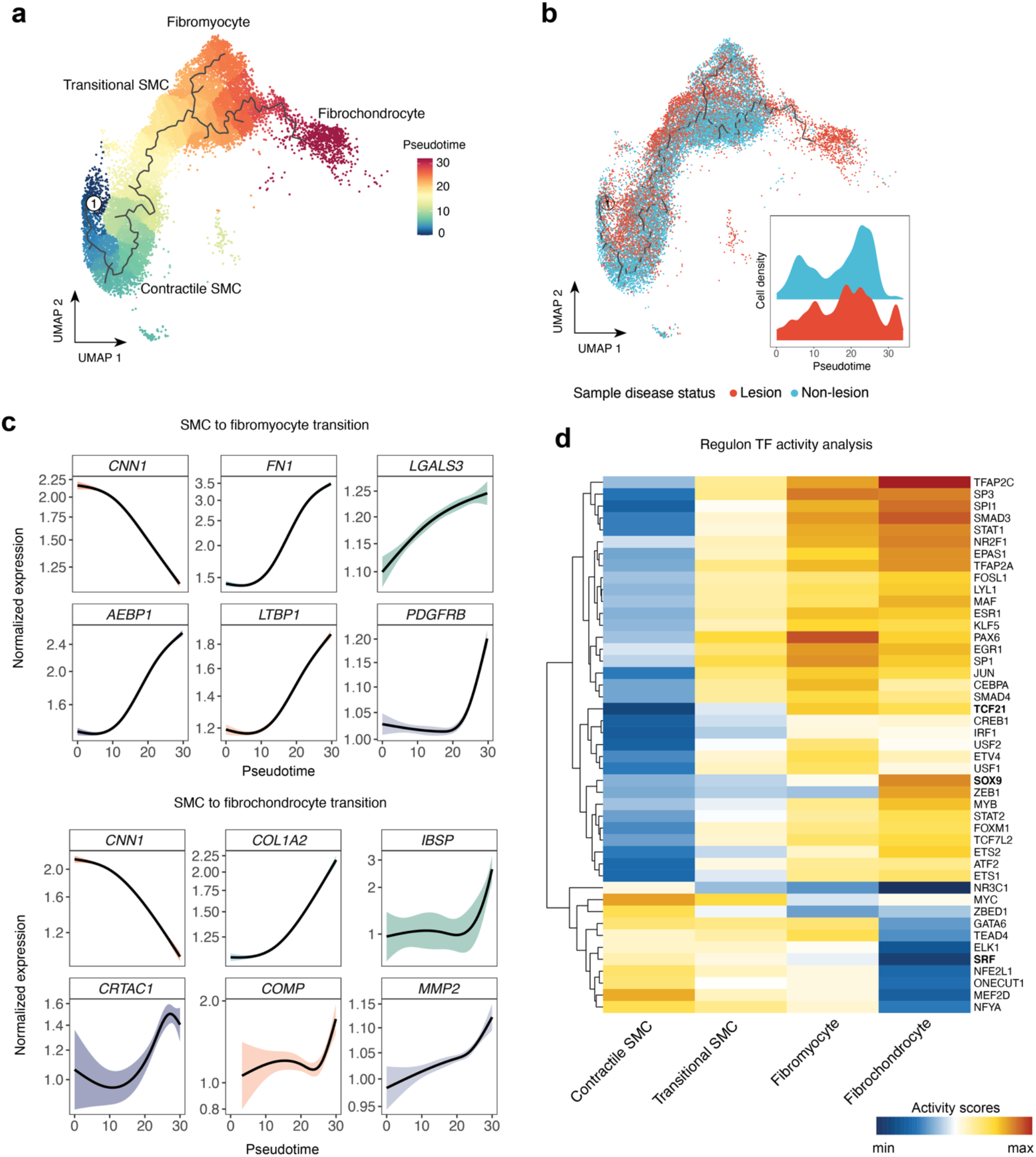
Pseudotime and TF inference activity for ECM-rich SMC phenotypes. (**a**) UMAP embeddings showing supervised pseudotime trajectory from Contractile to modulated SMCs calculated with Monocle 3^70^. SMC phenotypes for this analysis included contractile, transitional SMCs, fibromyocytes and fibrochondrocytes (FCs). The numbered circle depicts the root of the trajectory, which was defined as the subset of Contractile SMCs with highest *MYH11* expression. (**b**) Pseudotime trajectory with SMCs grouped according to lesion status. Inset plot depicts the density of cells from lesions and non-lesion libraries across pseudotime. (**c-d**) Cubic spline interpolation of SCTransform-normalized gene expression as a function of pseudotime. Genes plotted include hits from Monocle 3 and Seurat DE tests (FDR < 0.05). DE genes from SMCs to fibromyocytes trajectory: *FN1*, *LGALS3*, *AEBP1*, *LTBP1*, *PDGFRB.* DE genes from SMCs to FC trajectory: *COL1A2*, *IBSP*, *CRTAC1*, *COMP*, *MMP2*. (**e**) Transcription factor (TF) activity prediction with VIPER^74^ based on DoRothEA regulons^75^ for contractile and ECM-rich SMC phenotypes. Only regulons with confidence scores A-C (based on the number of supporting evidence) were used for this analysis. Highly variable TFs were selected for plotting and scale indicates relative predicted activity per TF.

Next, we modeled the expression dynamics of our DE genes and as expected, expression of canonical SMC contractile markers, *MYH11* and *CNN1* sharply decreased across pseudotime, whereas *ACTA2* and *TAGLN* persisted (**Fig 6c and Supplementary Fig. 6**). Interestingly, some fibromyocyte markers such as *FN1, AEBP1 and LTBP1,* in addition to the pioneer marker *LGALS3* showed a steady increase with adoption of the transitional state (**Fig. 6c**), while genes such as *PDGFRB,* were increased later suggesting a distinct role in the fibromyocyte state (**Fig. 6d**). In parallel, we inspected FC markers (**Supplementary Table 6**) and genes from module 9 and observed a steady increase in expression of *COL1A2* and *MMP2*, whereas *IBSP*, *CRTAC1* and *COMP* were increased at later pseudotime stages, presumably as transitional SMCs adopt a FC fate (**Fig. 6c, lower panel**).

### TF activity inference analysis

We next investigated the upstream transcription factors driving cell specific expression changes using TF activity inference with VIPER^74^ and the DoRothEA collection of well-curated and stable human regulons^75^. This analysis corroborated known regulators of fibromyocytes and FCs such as TCF21 and SOX9 (**Fig. 6d**). However, we also achieved granularity to detect changes in AP- 1 (e.g., JUN, FOSL), TEAD, ETV and ETS factors activity in fibromyocytes vs FCs. Interestingly, we also observed increased regulon activity of the TGF-β signaling mediator SMAD3 in fibromyocytes and FCs compared to contractile and transitional SMCs, with FCs showing the highest activity. To corroborate these results, we then interrogated our coronary artery snATAC-seq data^76^ using ArchR^77^ and besides confirming increased accessibility of AP1 factors, we found that accessible regions in the ECM-rich SMC cluster were specifically and highly enriched for SMAD3 motifs compared to contractile SMCs (**Supplementary Fig. 6**). This suggests that SMAD3 activity is critical as SMCs transdifferentiate towards more synthetic phenotypes.

### *CRTAC1 and LTBP1* as candidate markers of FC and fibromyocytes in human atherosclerosis

Fibromyocytes and FCs play major roles in atherosclerotic lesion stability^18, 20^, however, characterization of distinct markers for these SMC subtypes in humans has been limited. In addition to validating known markers of FCs (*CYTL1*, *COMP, COL1A1/2*)^22, 78^, we found that cartilage acidic protein (*CRTAC1*) was expressed 3-fold higher relative to other SMC clusters (**Fig. 4c, Supplementary Table 6**). *CRTAC1* is a specific marker of human chondrocytes during ossification^72, 79^, and is implicated in osteoarthritis^80^. However, it has not been identified in mouse atherosclerosis studies, potentially suggesting specificity for human SMCs. The top fibromyocyte markers (**Fig. 4c, Supplementary Table 6**) also revealed latent transforming growth factor binding protein 1 (*LTBP1*), a regulator of TGF-beta secretion and activation^81^, which may also contribute to SMC modulation and plaque stability.

Both *CRTAC1* and *LTBP1* are enriched in artery tissues compared to other tissues in GTEx and predominantly expressed in SMCs from level 1 annotations (**Supplementary Fig. 7**). We also observed expression of *CRTAC1* in SMCs enriched for the murine FC gene signature (**Fig. 4a and Fig. 7a**) along with the calcification marker, *IBSP*^20, 66, 78, 82^, while *LTBP1* showed gradual upregulation from contractile SMCs to fibromyocytes, consistent with our previous pseudotime results (**Fig. 6c**). Both *CRTAC1* and *LTBP1* were positively correlated with known osteochondrogenic or fibromyocyte markers, respectively (**Fig. 7b-c**), but *CRTAC1* was more negatively correlated with canonical SMC markers. This suggests expression of these genes in SMCs is associated with a progressive loss of the contractile phenotype and gain of synthetic and pro-calcification gene programs^83, 84^.

**Figure 7.**
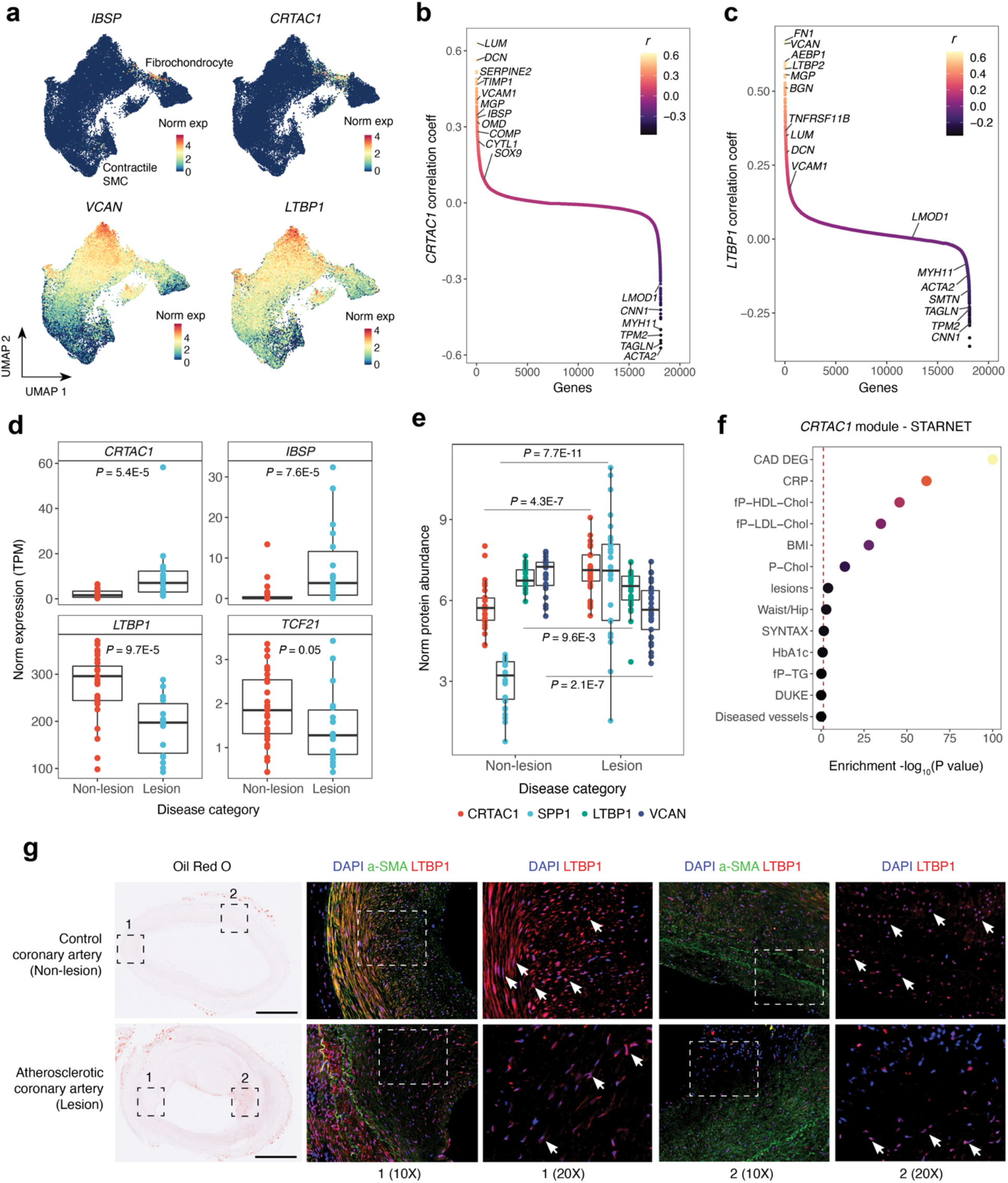
*CRTAC1* and *LTBP1* are novel candidate markers of atherosclerosis. progression. (**a**) UMAP embeddings for SCTransformed normalized expression of *CRTAC1* and *LTBP1* within the subset of the reference defined in 4a. *IBSP* is used as a control marker of calcification and *VCAN* as a control marker of fibromyocytes. (**b**) Pearson correlation plot of *CRTAC1* with every other gene across SMCs and FC clusters. (**c**) Pearson correlation plot of *LTBP1* with every other gene across SMCs and Fibromyocytes clusters. Selected examples of canonical contractile, ECM-related and osteochondrogenic genes regulated during SMC modulation are shown. (**d**) Bulk RNA-seq expression of *CRTAC1*, *IBSP, LTBP1* and *TCF21* in coronary arteries from lesion (n=27) and non-lesion samples (n=21). Data points represent normalized expression counts (TPMs). Shown *p* values were calculated using a non-parametric Wilcoxon Rank Sum Test **(Methods)**. (**e**) Log-normalized protein expression of calcification (CRTAC1, SPP1) and ECM-related (LTBP1, VCAN) genes in non-lesion (n=27) and lesion (n=29) samples. Disease category detailed in **Methods**. Shown *p* values were calculated using an unpaired Student’s *t*-test **(Methods)**. Boxplots in (**d**) and (**e**) represent the median and the inter-quartile (IQR) range with upper (75%) and lower (25%) quartiles shown, and each dot represents a separate individual. (**f**) Clinical trait enrichment for *CRTAC1*-containing module in a subclinical mammary artery in STARNET gene regulatory network datasets. Pearson’s correlation *p* values (gene-level) were aggregated for each co-expression module using a two-sided Fisher’s exact test. Case/control differential gene expression (DEG) enrichment was estimated by a hypergeometric test. (**g**) Representative Oil-red-O staining and immunofluorescence (IF) staining of LTBP1 in human atherosclerotic coronary artery segments from normal (control/subclinical, n=4) and stage IV-V lesions (n=4) (Stary classification). Two regions of interest per sample stained with LTBP1 are shown on the right, respectively at 10 and 20x magnification, with LTBP1 (red channel), alpha-smooth muscle actin (a-SMA; green channel), and nuclei stained with DAPI (blue). Arrowheads point to LTBP1-positive cells. Ruler =1mm.

To validate *CRTAC1 and LTBP1* as markers of human atherosclerosis, we queried in-house and public transcriptomic and proteomic datasets^85^. *CRTAC1* gene expression was significantly upregulated in advanced lesions compared to non-lesion control/subclinical samples (**Fig. 7d- e**). In contrast, both *LTBP1* mRNA and protein levels were downregulated in advanced coronary lesions, similar to known fibromyocyte genes *TCF21* and *VCAN* (**Fig. 7d-e, Supplementary Fig. 7**). Furthermore, *CRTAC1* was upregulated in unstable versus stable carotid plaque regions^85^, whereas *LTBP1* and other fibromyocyte genes showed the opposite trend. Next, we queried the Stockholm-Tartu Atherosclerosis Reverse Network Engineering Task (STARNET) gene regulatory networks across seven cardiometabolic tissues^86^. *CRTAC1* was identified as a significant key driver within its co-expression module (**Supplementary Table 11 and 12**), which was highly associated with CAD genes, C-reactive protein (CRP), and LDL cholesterol (**Fig. 7f**). The *CRTAC1* module was enriched for calcification related GO terms, whereas the *LTBP1* module was enriched for cholesterol traits and ECM remodeling terms (**Supplementary Tables 13-15, Supplementary Fig. 7**). This suggests *LTBP1* expression might be associated with an earlier stage of modulation, preceding calcification.

Immunofluorescence staining showed LTBP1 localization in medial and fibrous neo-intimal regions of early lesions, the expected location of fibromyocytes in control/subclinical and atherosclerotic coronary segments (**Fig. 7g and Supplementary Fig. 8**). We observed a more uniform neointimal expression in subclinical lesions compared to advanced lesions, which showed more heterogenous and reduced expression along the vulnerable and lipid-rich shoulder regions from representative samples. These data together support the potential role of LTBP1 as a marker of plaque stability, which could be further evaluated in genetic models of atherosclerosis or plaque rupture.

## Discussion

In this study we generate the first comprehensive single-cell transcriptomic atlas of human atherosclerosis, including 118,578 high-quality cells from atherosclerotic coronary and carotid arteries. We define cell subtypes which have not been previously identified from individual human atherosclerosis scRNA-seq studies. We leverage this scRNA-seq reference to investigate SMC phenotypic modulation in humans and identify etiologic SMC subtypes for GWAS cardiovascular diseases/traits, including CAD and CAC. We derive insights into SMC- myeloid cell crosstalk and gene regulatory networks in atherosclerotic lesions. Finally, we uncover two candidate SMC subtype-specific markers of plaque stability and calcification.

An inherent challenge in single-cell studies is labeling cell clusters, particularly when defining cell states or phenotypes. Our systematic labeling approach allowed corroboration of reported SMC murine phenotypes^18–20^, but also unveiled rare SMC clusters including a “foam-like” state, supporting previous studies reporting a SMC-derived foam-like phenotype upon exposure to lipoproteins^87, 88^ and in human lesions^89^. The lower abundant foam-like SMCs in individual reports might be explained by tissue digestion and single-cell isolation procedures. Cells in this cluster expressed lipid metabolism genes (e.g., *APOE*, *APOC1)* but not other traditional foamy macrophage markers, in line with previous findings^89^. Expression of ECM genes such as *TIMP1* suggest SMC-derived foam cells may acquire a unique gene signature. This aligns with evidence suggesting that modulated SMCs accumulate lipids in plaques without adopting monocyte-derived foam macrophage transcriptional signatures^18^.

Our granular SMC annotations also defined etiologic SMC phenotypes for cardiovascular diseases and traits. Previous work from our group and others^90–92^ established a substantial contribution of SMCs towards CAD risk. We further deconvolved the SMC signal, prioritizing fibromyocytes and foam-like SMCs underlying cardiovascular diseases. We demonstrate enrichment of a fibromyocyte-specific transcriptional signature in CAD and MI heritability, supporting their role in plaque stability, and the importance of targeting this SMC phenotype for future therapeutic interventions. These heritability analyses also linked SMC-derived FCs to CAC, an established pathological hallmark of subclinical and advanced atherosclerotic lesions^93^.

Though fibromyocytes and FCs originate from SMCs^18–20, 94^, these two ECM-producing phenotypes play distinct roles in plaque stability^9^ and may follow shared or unique fates. Using pseudotime, we revealed a branchpoint where transitional SMCs could adopt either fibromyocyte or FC fates. This, however, does not preclude the possibility that fibromyocytes could be primed to become FCs as *Cheng et al*^67^ and our TF activity inference results suggest, but implies an alternative modulation path. However, additional lineage-tracing experiments are needed to address this possibility. To refine the SMC regulatory landscape, we validated known vascular TFs such as TCF21 and SOX9, but also mitogenic TFs such as MYC in transitional SMCs and AP-1 and TEAD factors across fibromyocytes and FCs. Interestingly, we also observed higher SMAD3 motif accessibility in ECM-rich SMCs (fibromyocytes/FCs) compared to contractile SMCs in our snATAC data^76^. This CAD causal gene^90, 95^ is known to stimulate chondrogenesis in mesenchymal stem cells by enhancing Sox9 transcriptional activity^96, 97^ and previous reports reveal Smad3 activity in murine FCs^78^. Consistently, our TF regulon analysis confirmed increased SMAD3 regulatory activity along the transitional SMC-fibromyocyte-FC continuum. We thus speculate that SMAD3 plays different roles in modulated SMC phenotypes, and its association with CAD risk might be at least in part related to its activity in FCs.

Our work also provides further insights into fibromyocyte and fibrochondrocyte (FC) gene signatures in human lesions. We identified *CRTAC1* as a top, previously uncharacterized FC marker with increased expression observed in coronary lesions. While *CRTAC1* was also upregulated in unstable carotid plaques^85^, its role in plaque rupture is unknown. More exhaustive *ex vivo* and *in vivo* functional characterization is required to assess its role in diverse calcification phenotypes^98–100^ and plaque stages. Our analyses also revealed *LTBP1* as a specific fibromyocyte gene. Previous quantitative trait loci (QTL) studies have shown that the risk allele of CAD SNP rs6739323 is associated with lower LTBP1 expression in cultured human SMCs^101^, and human aortic tissue^102^. Here, we confirmed *LTBP1* expression in native contractile SMCs in the medial coronary layer, and in fibromyocytes in the fibrous neointima. Elevated *LTBP1* expression in stable carotid plaques^85^ or subclinical coronary arteries suggests a protective role promoted by transitioning SMCs towards fibromyocytes. Though *CRTAC1* and *LTBP1* show promise as disease biomarkers, follow-up experimental studies are necessary to confirm their precise functions in atherosclerosis.

We acknowledge the potential over-simplification of our lesion status in our group comparisons. First, the non-lesion samples were extracted from patients with non-ischemic dilated cardiomyopathies (DCM) without discernible atherosclerosis^17^. The higher representation of inflammatory macrophages in these libraries could be a consequence of myocardial inflammation or secondary subclinical diffuse intimal thickening. While most of the cell types were balanced across samples, we observed under-represented fibroblasts in one of the carotid lesion datasets^19^, most likely due to the sample preparation. Finally, the lesion definition in human samples is inherently less refined both spatially and temporally compared to murine studies. Nevertheless, our atlas represents a valuable integrated resource of the most relevant arterial beds to atherosclerosis and provides a comprehensive map of cell diversity in human lesions (**Supplementary Fig. 9**). With newly generated single-cell datasets, there will be a need to address the variability of reported phenotypes and create a unified map of atherosclerosis.

Integrative single-cell meta-analyses are a powerful discovery tool to capture both robust and subtle signals and are expected to improve with future iterations. Ultimately this will catalyze mechanistic and translational studies and contribute towards developing novel therapeutic strategies for complex diseases such as CAD.

## Supplementary Figures

**Supplementary Figure 1.**
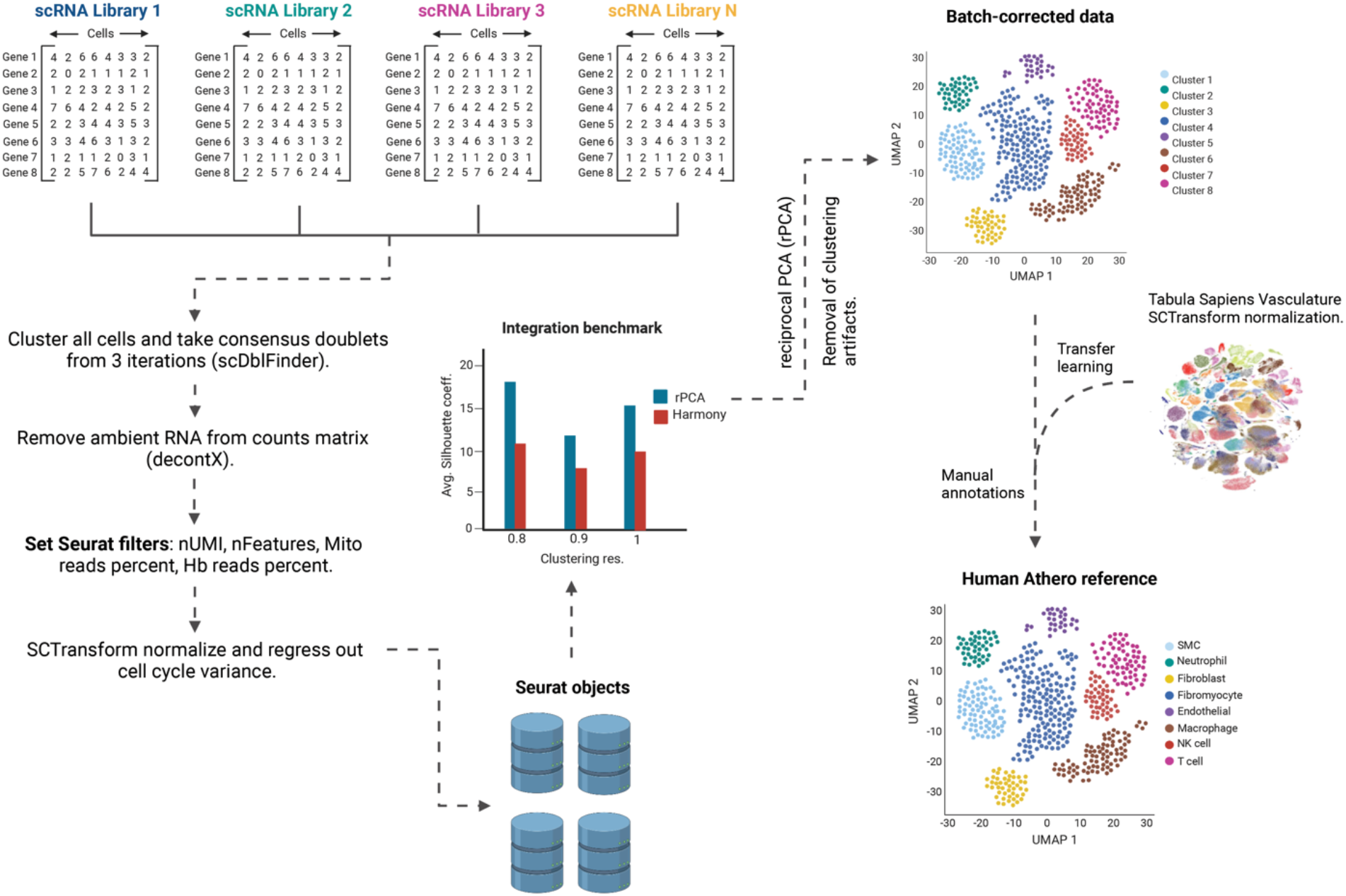
Pipeline used to build the integrated scRNA reference. Workflow for standardizing processing of each scRNA-seq library and integration. Doublets were first identified and removed using scDblFinder^104^. Upon doublet removal, ambient RNA was removed using decontX^25^. The decontaminated matrix was then used for downstream filtering of cells based on 1) number of detected genes 2) number of UMIs 3) percentage of reads mapping to mitochondrial genome 4) percentage of reads mapping to hemoglobin genes using Seurat^26^. Libraries were normalized using SCTransform^27^ integrated using four approaches (CCA + MNN, rPCA^26^, Harmony^32^, Scanorama ^31^). PCA embeddings from each approach were then used for measuring inverse local Simpson index (LISI) scores and silhouette coefficients. rPCA was used to harmonize the 22 included sequencing libraries and level 1 labels were added using Transfer learning with the Tabula Sapiens Vasculature subset as reference. Finally, a combination of automated^48^ and manual curation was carried out to identify more granular cell subtypes (level 2).

**Supplementary Figure 2.**
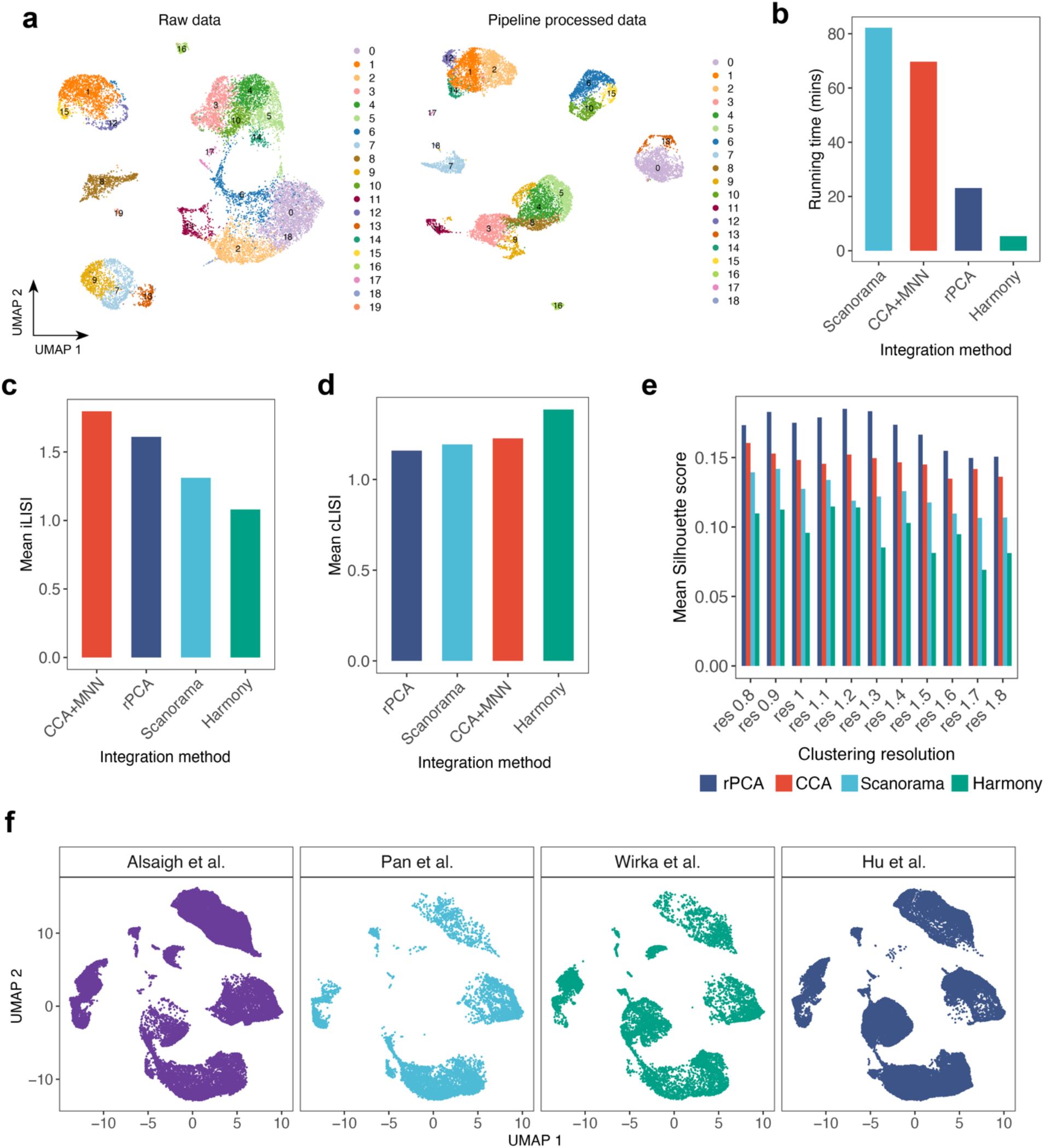
scRNA pipeline QC and integration benchmark metrics. (**a**) Example UMAP embeddings of one of the included libraries before and after going through our scRNA-seq processing pipeline. (**b**) Running time of each of the four integration approaches tested. The Y axis shows time in minutes. (**c**) Mean integration LISI (iLISI) scores calculated for each integration approach. Higher iLISI scores depict improved mixing and batch removal. (**d**) Mean cell type LISI (cLISI) calculated for each integration approach. Lower cLISI scores represent increased biological conservation. (**e**) Mean silhouette coefficients calculated for each integration approach. Silhouette coefficients were calculated using euclidean distances across a range of clustering resolutions (0.8-1.8) to determine optimal clustering resolutions. Silhouette scores range from (-1, 1) where higher scores depict improved clustering quality or purity. PCA embeddings (30 PCs) were used for calculation of metrics in (**c**-**e**). (**f**) UMAP embeddings of the integrated reference showing the contribution of the included studies to each cell compartment, confirming proper removal of batch effects. For additional details on processing and benchmark see **Methods**.

**Supplementary Figure 3.**
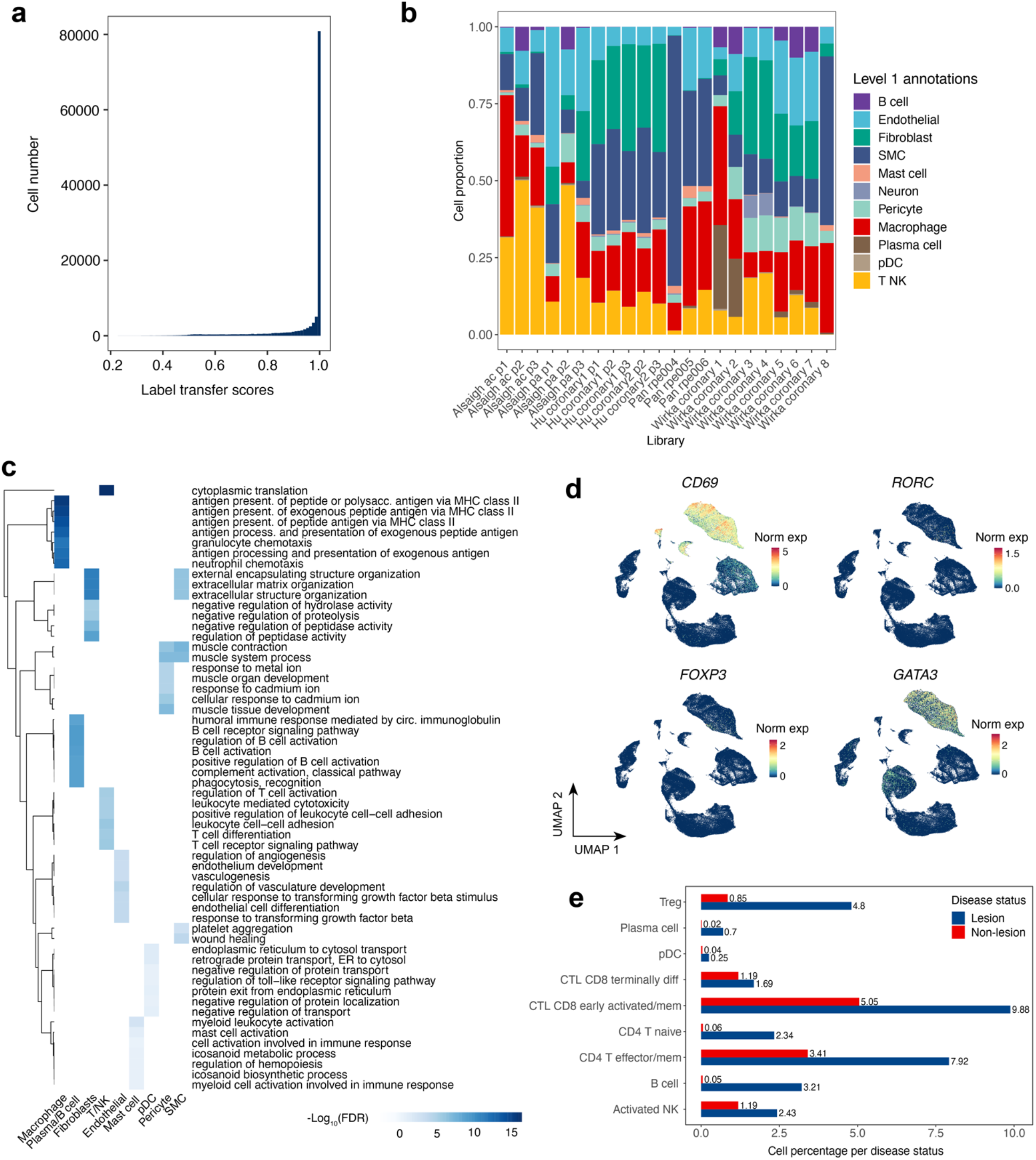
Additional characterization of level 1 and level 2 cell type annotations. (**a**) Confidence scores from predicted labels using the Seurat Transfer learning classifier with the TS vasculature reference. Confidence scores range from 0-1 where higher scores refer to unambiguous cell label calls. (**b**) Bar plot showing the distribution of level 1 annotated cell types across the 22 sequencing libraries included in this study. (**c**) Gene set enrichment analysis (GSEA) for level 1 annotated cell types. This analysis was carried out with gProfiler2^105^ and the top seven significantly enriched terms (FDR <0.05) were selected for plotting. (**d**) UMAPs of SCTransform-normalized expression of genes defining key T cell states and subtypes (*CD69*: early activation; *RORC*: Th17 cells; *FOXP3*: Regulatory T cells (Treg); *GATA3*: Th2 cells. (**e**) Bar plot of level 2 Lymphoid cell subtypes according to lesion status, in percent. Frequencies are normalized to the total number of cells, respectively (lesion n=59691; non-lesion n=58887).

**Supplementary Figure 4.**
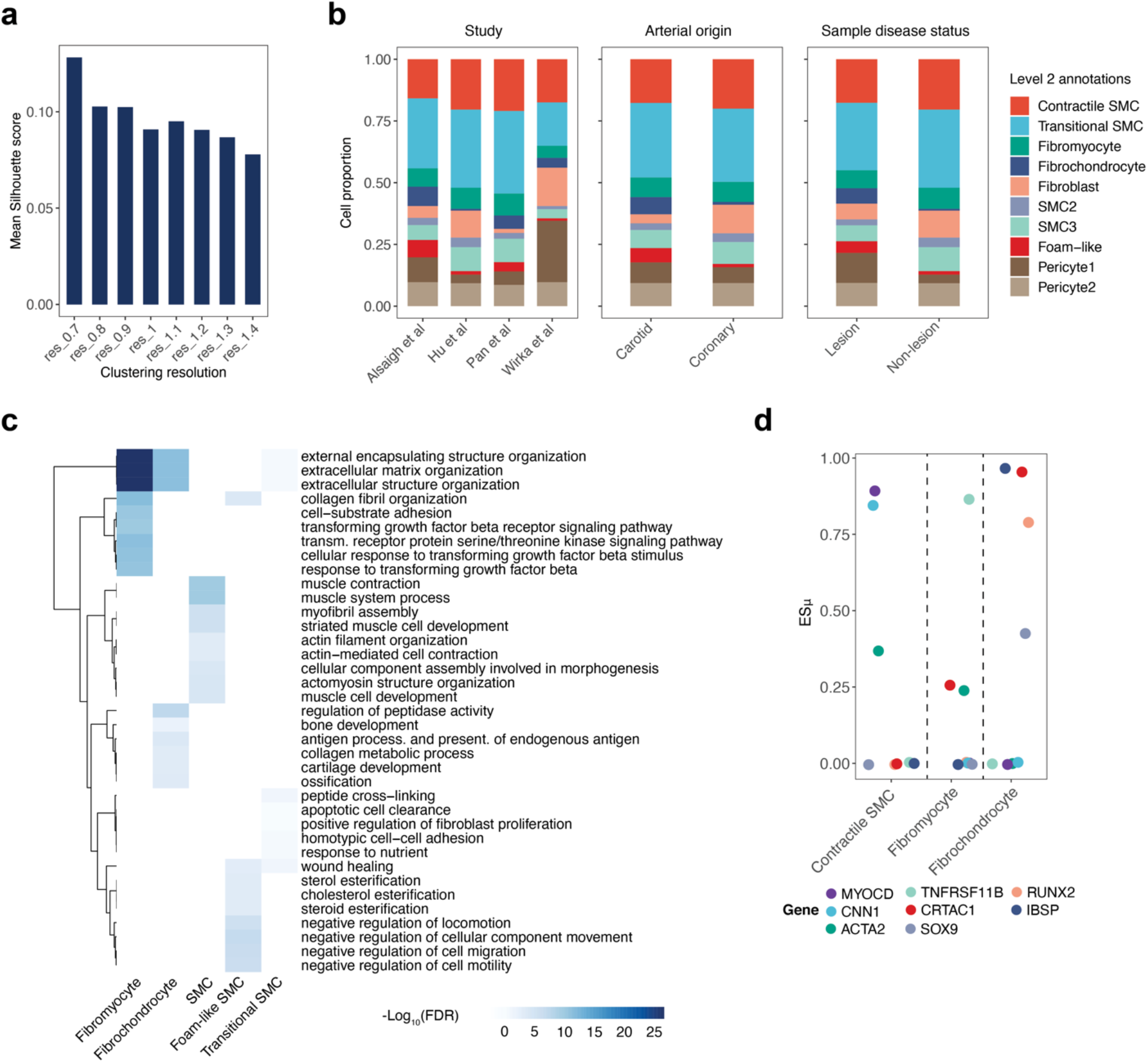
Additional characterization of SMC subtypes. (**a**) Barplot depicting silhouette analysis using PCA embeddings for reference subset (SMCs, Pericytes and Fibroblasts). Silhouette coefficients are calculated across a range of resolutions to find optimal parameters for subclustering and range from (-1, 1) where higher scores depict improved clustering quality or purity. (**b**) Stacked bar plot of SMC level 2 annotations, as pericytes and fibroblasts across studies, arterial beds (Coronary or Carotid) and sample disease status (lesion and non-lesion). (**c**) Dot plot showing ESμ values for canonical contractile markers (*MYOCD*, *CNN1*, *ACTA2*) as well as synthetic (*TNFRSF11B*) and osteochondrogenic markers (*RUNX2*, *SOX9*, *IBSP*) of SMC modulation for Contractile and ECM-rich SMC phenotypes (Fibromyocytes and FCs). ESμ values were plotted from a gene expression specificity matrix generated with CELLEX^103^. For additional details on ESμ values see **Methods**. (**d**) Gene set enrichment analysis (GSEA) for level 2 annotated SMCs. This analysis was carried out with gProfiler2^105^ and the top nine significantly enriched terms (FDR < 0.05) were selected for plotting.

**Supplementary Figure 5.**
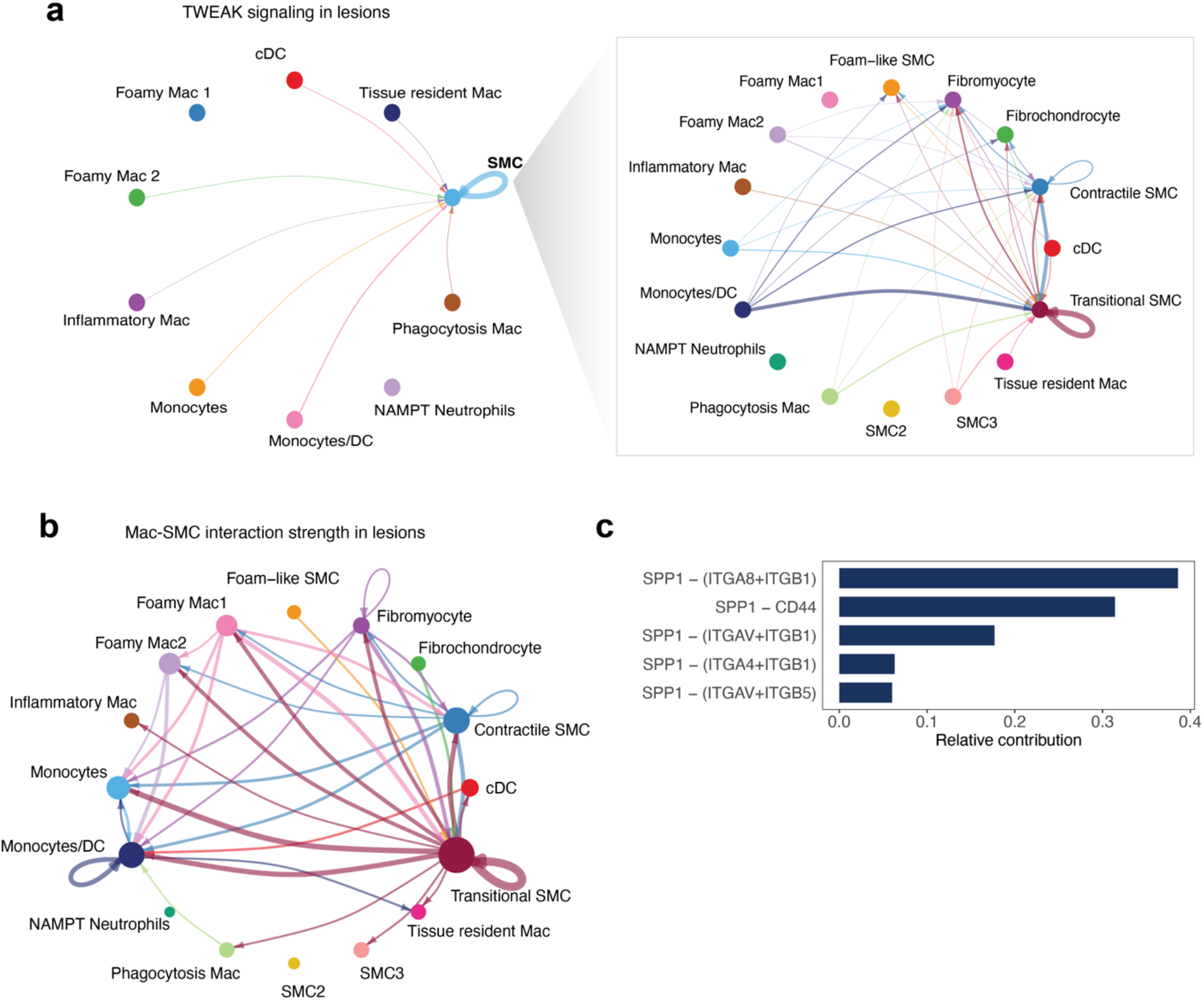
TWEAK and SPP1 signaling for myeloid and SMC cell types. (**a**) Circle plot depicting sources and targets for TWEAK signaling using level 2 labels for myeloid cells and level 1 SMC labels (left) and using level 2 labels for myeloid and SMCs (right) Width of the edges depicts weight or strength of the interaction based on calculated communication probability between a pair of cell types. Interactions were calculated using the CellChat^69^ human database. (**b**) Circle plot showing the aggregated cell-cell communication network for level 2 Myeloid and SMC labels. Top 15% of interactions are shown in the plot. (**c**) Bar plot showing the relative contribution of each ligand-receptor pair for SPP1 signaling. Width of the edges in the circle plot depicts the weight/strength of the interactions in **a**,**b**.

**Supplementary Fig. 6.**
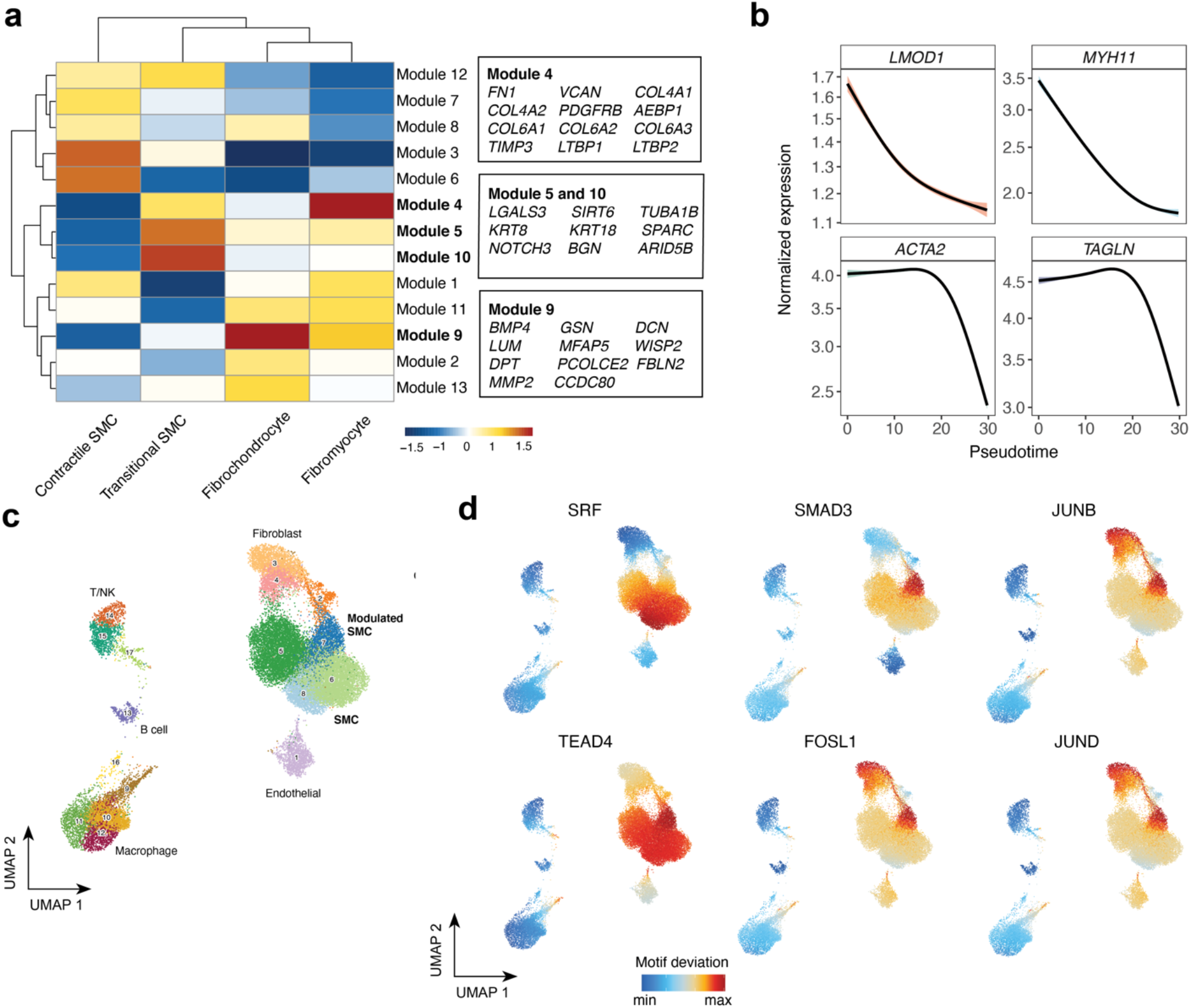
SMC gene expression dynamics through pseudotime and snATAC-seq TF activity inference. (**a**) Heatmap of varying gene module expression as Contractile SMCs transition into ECM-rich phenotypes (fibromyocytes and FCs). Differential genes across pseudotime were calculated using graph autocorrelation analysis with Monocle 3 and then grouped into modules using Louvain community analysis. Color scale represents aggregated expression of genes in each module across the above-mentioned SMC phenotypes. Boxes (right) list key genes found in each module. (**b**) Cubic spline interpolation of SCTransform-normalized expression of canonical contractile markers (*LMOD1*, *MYH11*, *ACTA2*, *TAGLN*) as a function of pseudotime. (**c)** UMAP and Louvain clustering of coronary arteries snATAC-seq data. Each dot represents an individual cell colored by cluster assignment. Cell type labels in bold represent Contractile and ECM-rich modulated SMC populations as defined in *Turner et al*^76^. (**d**) UMAPs of ChromVAR TF motif accessibility deviation scores for factors shown as highly variable in previous SMC analysis with DoRothEA regulons (SRF, TEAD4, SMAD3, FOSL1, JUN).

**Supplementary Figure 7.**
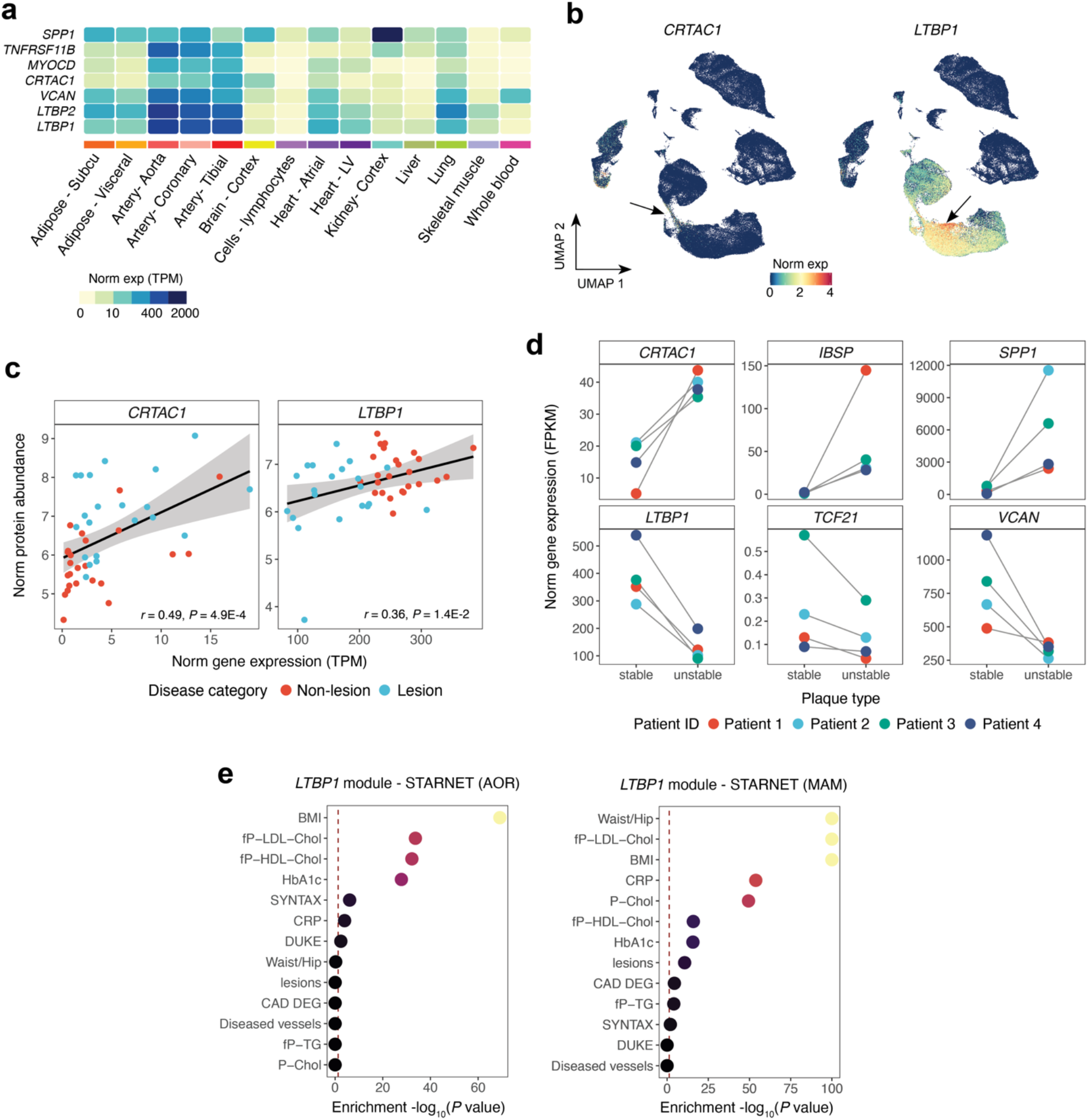
Bulk and single-cell validation for *CRTAC1* and *LTBP1* activity across atherosclerosis-relevant arterial beds. (**a**) Heatmap showing normalized expression (TPMs) of canonical contractile, ECM-related and calcification genes expression across several GTEx tissues. (**b**) UMAP plots of *CRTAC1* and *LTBP1* across the entire integrated reference. SCTransform-normalized gene expression is indicated by color. (**c**) Scatter plot showing correlations between RNA and protein levels for matching lesion and non-lesion coronary artery samples (n=42 samples). Pearson correlation coefficients and corresponding *p* values were calculated and included within each plot **(Methods)**. (**d**) Dot plot showing normalized expression (FPKMs) for calcification-related genes (*CRTAC1*, *IBSP*, *SPP1*) and ECM-related genes (*LTBP1*, *TCF21*, *VCAN*) from a public RNA-seq dataset of human fresh carotid lesions^85^. Dots of the same color represent matched patient (n=4) samples of stable and unstable plaque regions (stable, n=4; unstable, n=4). (**e**) Clinical trait enrichment for *LTBP1*-containing module in aorta and subclinical mammary arteries in STARNET gene regulatory network datasets. Pearson’s correlation *p-*values (gene-level) were aggregated for each co-expression module using a two-sided Fisher’s exact test. Case/control differential gene expression (DEG) enrichment was estimated by a hypergeometric test.

**Supplementary Figure 8.**
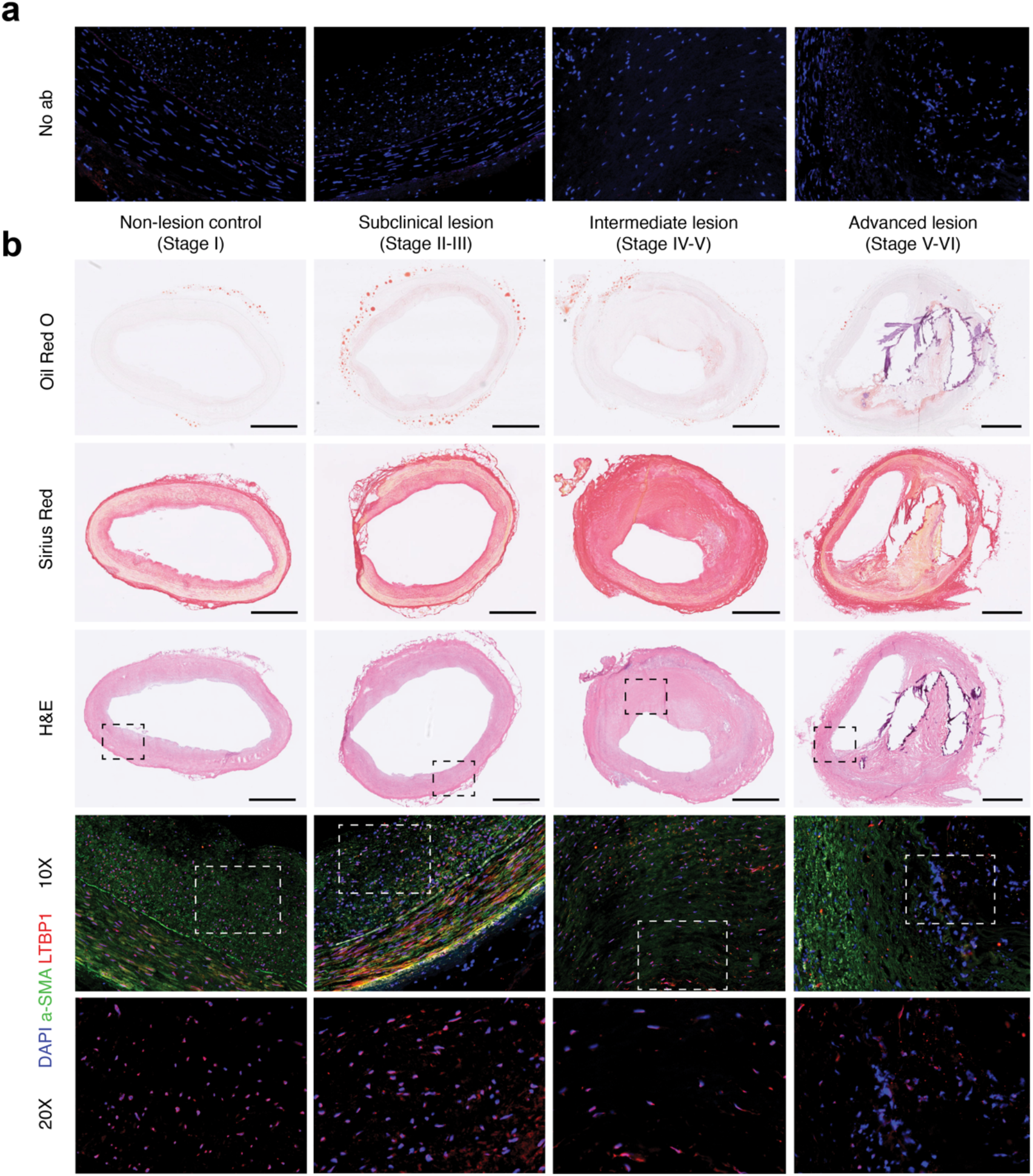
Histology and immunostaining for LTBP1 protein in coronary atherosclerotic sections. (**a**) Representative negative controls (no primary antibody) immunofluorescence (IF) staining in human coronary arteries spanning controls, and stage II-VI lesions. Similar results were observed across n=8 independent donors. (**b**) Oil-red-O, Sirius Red and hematoxylin & eosin (H&E) histology staining of distinct human coronary artery segments from control-stage II, stage II-III, stage IV-V and stage V-VI lesions based on Stary classification stages. Regions of interest (ROIs) are showed on the H&E images from each sample for visualization with immunostaining at x10 and x20 magnifications. (**c**) Representative IF staining in the ROIs defined in b of corresponding human coronary artery samples with LTBP1 (red). Alpha-smooth muscle actin (a-SMA; green) and nuclei counterstained with DAPI (blue). Staining shown is representative of n=8 independent donors. Negative control images in (**a**) show matching region to that of LTBP1-stained samples in (**c)** carried out in adjacent tissue sections. White arrows highlight LTBP1/DAPI positive cells. Ruler =1mm.

**Supplementary Figure 9.**
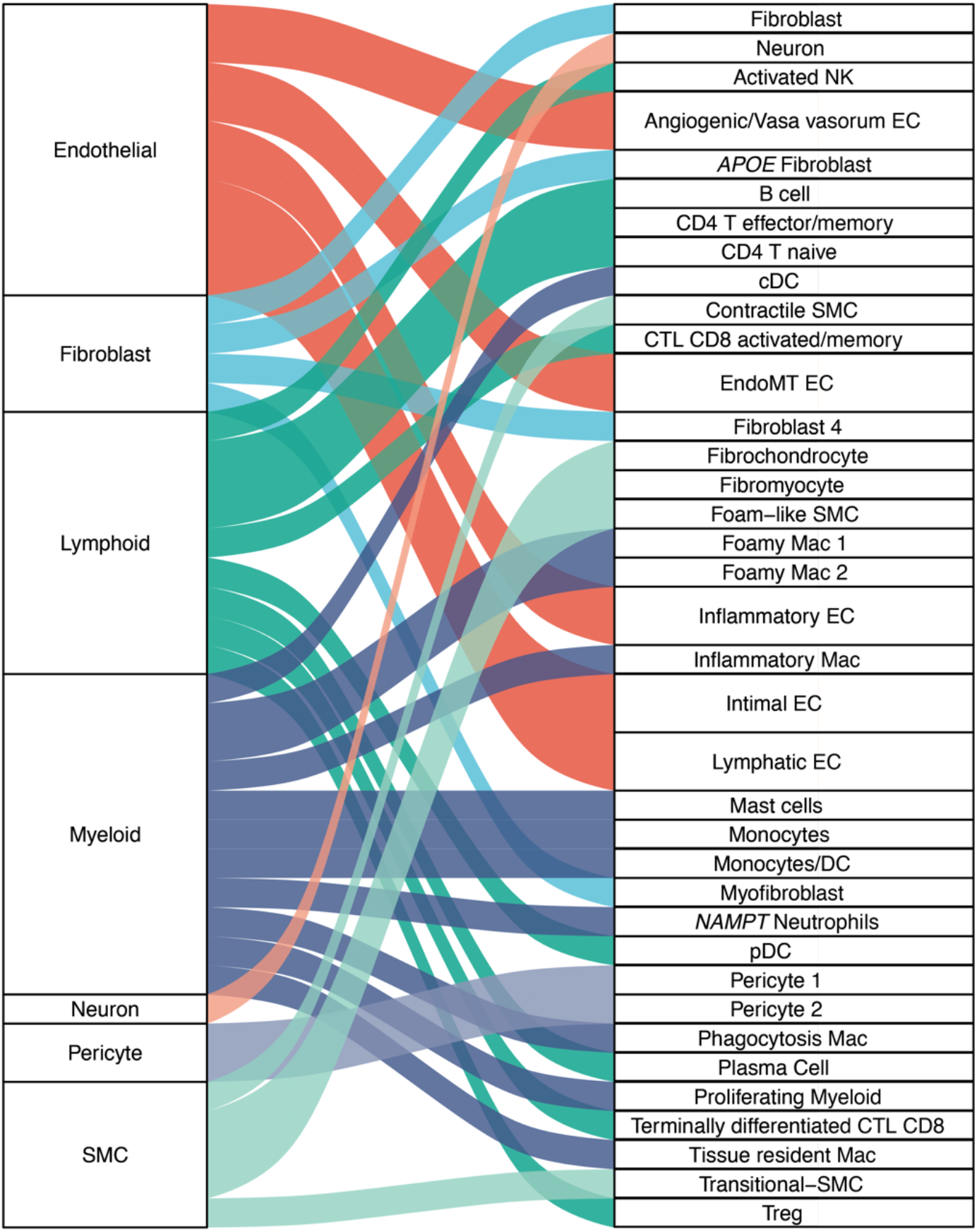
Summary of cell type diversity in human atherosclerosis. Riverplot depicting the relationship between level 1 cell compartments and level 2 cell subtypes for vascular and immune lineages. This plot was generated using the ggalluvial R package.

## Online Methods

### Ethics statement

Details regarding data collection for scRNA-samples included in this meta-analysis can be found in each publication^16–19^. Collection of coronary artery samples for bulk RNA-seq/proteomics data generation as well as histology and immunofluorescence analyses described in this manuscript complies with ethical guidelines for human subjects research under approved Institutional Review Board (IRB) protocols at Stanford University (no. 4237 and no. 11925) and the University of Virginia (no. 20008), for the procurement and use of human tissues and information, respectively.

### QC and normalization of scRNA-seq sequencing libraries

Raw count matrices from each library across the 4 studies were downloaded from GEO and Zenodo **(Data Availability, Supplementary Table 1)**. Processing for each of the 22 sequencing libraries was standardized in the following manner: Each library was loaded into the R programming environment (v.4.0.3) using Seurat^26^ (v.4.1.0). For each library we did a first pass of clustering with SCTransform normalization^27^ without removing low-quality cells.

In order to remove doublets, we referred to a recent benchmark of doublet-removal tools^106^ and chose the scDblFinder R package^104^ (v.1.4.0) given its superior accuracy compared to other tools. Seurat objects for each library were converted to SingleCellExperiment objects and used as input to generate artificial doublets using the cluster-based modality of scDblFinder. Briefly, scDblFinder creates a K-Nearest Neighbors graph using the union of real cells and artificial doublets and estimates the density of artificial doublets in the neighborhood of each cell. Since artificial-doublet generation approaches tend to display slight variance across different runs, we only kept consensus doublets from 3 iterations of the above-described process. Cell-barcodes that were marked as doublets were then removed from each raw counts matrix.

Ambient RNA contamination is a key issue during 10x protocols and can negatively impact clustering and extraction of gene markers. To filter out reads from ambient RNA, we ran DecontX^25^ within the celda R package (v1.6.1) in doublets-filtered raw counts matrices using default parameters. The decontaminated raw count matrices output by DecontX were then added into each Seurat object. We then set quality filters to keep cells that had 1) >= 200 and <= 4000 uniquely expressed genes 2) >= 200 and <= 20000 UMIs 3) <= 10% of reads mapped to the mitochondrial genome; cells with high percentages of reads mapped to mitochondrial genomes are considered to be low quality as this indicates cell membrane breaches and 4) <= 5% of reads mapped to hemoglobin genes since these cells likely depict contaminating erythrocytes as done in *Alencar et al*.

Raw count matrices were then normalized using SCTransform^27^ with parameter (vst.flavor=v2), which accounts for sequencing depth variability across cells. This omits the need for heuristic steps such as log-transformation and it has been shown to improve variable gene selection, dimensionality reduction and differential expression^27^. To avoid clustering results confounded by cell cycle state, cell cycle variance was regressed out during SCTransform normalization. We then carried out dimensionality reduction of the normalized counts matrix using PCA. The first 30 principal components (PCs) were used as input for clustering in Seurat, which relies on a Shared-Nearest-Neighbors (SNN) and Louvain community detection approach. We then applied Uniform Manifold Approximation and Projection (UMAP) non-linear dimensionality reduction using the first 30 PCs. UMAP embeddings were used for visualization of Louvain clustering results. Processed matrices were then stored as seurat objects for batch-correction.

### Integration benchmarking and building the reference

The choice of single-cell integration approach highly depends on the context of the individual datasets, the magnitude of batch effect and cell number. In order to harmonize processed sequencing libraries, we selected the following methods recommended from three recent benchmarks^28, 29, 107^ of single-cell transcriptomic data integration: Canonical Correlation Analysis + Mutual Nearest Neighbors (CCA + MNN), reciprocal PCA (rPCA)^26^ (v.4.1.0), Harmony^32^ (v.1.0) and Scanorama^31^ (v.1.7.1). We focused on four different metrics to choose a method: running time, efficiency of batch effect removal as denoted by the integrative Inverse Local Simpson Index (iLISI), conservation of biological variation using the “cell type” LISI (cLISI) and clustering purity measured by silhouette coefficients. The silhouette provides a measure of how well each cell has been classified by measuring how similar it is to its own cluster (cohesion) compared to other clusters (separation). For the benchmark, we used a subset of the data including 3 studies: *Wirka et al*, *Alsaigh et al* and *Pan et al*. Libraries from these studies were integrated as follows:

#### CCA + MNN

we created a list of selected Seurat objects and then selected 3000 highly variable genes. Integration with those variable genes was done using the PrepSCTIntegration(), FindIntegrationAnchors() and IntegrateData() functions. The batch-corrected expression matrix was then used for PCA dimensionality reduction, creation of the shared-nearest-neighbors (SNN) graph using 30 PCs and visualization with UMAP embeddings.

#### Harmony

libraries were first stored into a list and highly variable genes extracted using the function SelectIntegrationFeatures(). Libraries were merged into a single seurat object, and the list of highly variable genes was used for PCA dimensionality reduction. We used the first 30 PCs as input for RunHarmony() from the harmony package (v1.0), setting sequencing libraries (sample column in metadata) as the variables to correct for batch effects. Harmony embeddings were used for subsequent generation of the SNN graph, Louvain clustering and visualization with UMAP by setting reduction=”Harmony” within the FindNeighbors() and RunUMAP() Seurat functions and using the first 30 PCs.

#### rPCA

we created a list of processed Seurat objects and extracted the 3000 most highly variable genes using SelectIntegrationFeatures(). We then ran PCA across each library using the 3000 variable genes, identified integration anchors using FindIntegrationAnchors() setting reduction=”rpca” and harmonized datasets using IntegrateData(). As done for CCA, the batch-corrected expression matrix was then used for PCA dimensionality reduction, creation of the shared-nearest-neighbors (SNN) graph using 30 PCs and Louvain clustering followed by visualization with UMAP embeddings.

#### Scanorama

We used the reticulate R package (v.1.18) to import the Scanorama python module (v.1.7.1) into the R environment. We created a list with seurat objects containing the datasets to be integrated and stored normalized SCTransform-normalized counts and gene names for each dataset into a new list. We then batch corrected the data using the function using the correct() function from the Scanorama package setting the following parameters (return_dimred=TRUE and reurn_dense=TRUE). The batch-corrected expression matrix output by correct() was used to create a new Seurat object and Scanorama-produced dimensionality reduced embeddings were inserted into the Seurat object using the CreateDimReducObject() function. Scanorama embeddings were subsequently used to create a shared-nearest-neighbors (SNN) graph for Louvain clustering and for visualization with UMAP using the first 30 PCs.

#### Running time measurements

Running times for each integration task were then measured using base R Sys.time() functions. Sys.time() was defined at the beginning and the end of each integration task and then the time difference was calculated as end_time - start_time.

#### Silhouette analysis

Here we measured the quality or “goodness” of resulting clusters using the silhouette coefficient. For silhouette analyses, we extracted PCA embeddings from seurat objects with CCA+MNN, rPCA, Harmony and Scanorama integration outputs keeping the first 30 PCs. We then used these embeddings to compute an Euclidean distance matrix. Cluster IDs for each cell were obtained iteratively across a range of clustering resolutions (0.8-1.8) and Euclidean distance matrices were used to calculate silhouette width values using the cluster R package (v.2.1.0). The purpose of using the above range was to control for the clustering granularity parameter and to identify a range of clustering resolutions that would not lead to over- or underclustering of the data.

#### Calculation of LISI scores

Briefly, iLISI scores are a measure of the diversity within each cell neighborhood on a K-nearest-neighbor (KNN) graph. Higher iLISI scores depict increased mixing of batches within a cell neighborhood and therefore suggest improved removal of batch effects. For each of the integration methods described above we extracted PCA embeddings (30 first PCs) from the corresponding integrated Seurat object. We then created a data frame with each row corresponding to one cell and columns depicting batch variables (“Study”). We then computed iLISI scores for each cell using the compute_lisi() function from the lisi R package^32^ (v.1.0). Mean iLISI values were plotted and compared across different integration methods. cLISI scores, in turn, are considered a metric that measures conservation of biological variation. With the assumption that each cluster should generally harbor cells from the same type, we created a dataframe with each row corresponding to each cell and a column depicting Louvain cluster identities. cLISI scores for each cell were calculated and plotted as described above.

#### Integration of scRNA libraries and additional quality control

Upon determining the appropriate integration approach for the datasets of interest, we used rPCA to harmonize the 22 sequencing libraries as described in the above section. Upon integrating libraries, we reduced dimensionality of the data using PCA. A SNN graph was constructed using 50 nearest neighbors and the first 30 PCs as input. Clusters were identified using the above graph with a resolution of 1, which was within the range of higher mean silhouette coefficients from the previous benchmark. Gene markers for each cluster were identified using PrepSCTMarkers() and the Wilcoxon Rank Sum test as implemented in the FindAllMarkers() function from Seurat (v.4.0). We considered genes that were expressed in at least 25% of the clusters being compared (one cluster vs all others) and that had a logfc.threshold=0.25. Genes fulfilling that criteria in addition to having multiple-testing adjusted P-values <= 0.05 were considered as differential cluster markers. Upon inspection of the gene signatures of each cluster, we found two small clusters comprising 432 cells (0.36% of cells in the integrated reference) expressing markers of multiple major lineages, which likely represent residual doublets and thus were removed from the reference. Upon removing these residual doublets, cells were re-clustered using the above-described parameters. Finally, inspection of cluster markers expression in UMAP space allowed us to identify and remove UMAP artifacts (e.g., cells with Natural Killer signatures within candidate fibroblast clusters). The 306 cells (0.25% of cells in the reference) comprising these artifacts were removed to obtain the final iteration of the reference. This step was necessary to ensure robustness of cell type and subtype annotations as well as other downstream analyses.

### Cell type annotations

To annotate cell types in the integrated reference, we used a systematic approach to define broad labels (level 1) as well as more granular cell subtype labels (level 2).

#### Level 1 annotations

To define broad cell type partitions, we accessed public data from the Tabula Sapiens (TS) consortium (https://tabula-sapiens-portal.ds.czbiohub.org/organs). To improve the specificity of annotations, we downloaded the vasculature subset of this transcriptomic atlas. Upon downloading the TS vaculature h5ad file, this dataset was converted into a Seurat-compatible format using the SeuratDisk R package (v.0.0.0.9019). To match the normalization workflow described in the scRNA sequencing library processing section, we extracted the TS vasculature raw counts matrix and normalized gene expression data using SCTransform. We then applied Seurat’s reference-based transfer learning (using FindIntegrationAnchors() and TransferData() to annotate cells in our meta-analyzed reference. In this case, the TS vasculature seurat object with author-provided cell type annotations was defined as reference for label transfer. Confidence scores of predicted labels ranging from 0-1 (where 1 indicates that labels were annotated in a fully unambiguous manner) were extracted from the output of TransferData() and are shown in the Data Supplement. Gene markers for level 1 annotations were obtained using the PrepSCTMarkers() and FindAllMarkers() functions from Seurat (v.4.1.0) setting the following thresholds: logFC=0.25 and min.pct=0.25

#### Level 2 annotations for endothelial, fibroblasts and immune cells

To define more granular cell subtypes for the meta-analyzed data, we used a combination of automated and manual annotations. We first annotated cell subtypes for endothelial, myeloid and lymphoid lineages using markers from atherosclerosis murine scRNA meta-analyses of SMCs and immune cells as well as relevant human atherosclerosis scRNA studies^21–23, 42–47^. Annotations using curated markers from the literature were corroborated with the assistance of experts at UVA. To further confirm and inspect immune cell subtype annotations in our reference, we used a logistic-regression with stochastic^48^ gradient-descent framework implemented by the command-line tool CellTypist^48^. CellTypist leverages a database of 20 different tissues and 19 reference datasets with a focus on myeloid and lymphoid cells. Specifically, we applied CellTypist low-hierarchy classifiers (using the Immune_All_Low.pkl and Immune_All_AddPIP.pkl models which harbor 90 and 101 cell types, respectively) to our SCT-normalized reference counts matrix using both default settings as well as the majority voting classifier. Gene markers for level 2 annotations were obtained using the PrepSCTMarkers() and FindAllMarkers() functions from Seurat (v.4.1.0) setting the following thresholds: logFC=0.25 and min.pct=0.25

#### Level 2 annotations for SMCs

To explore SMC diversity in human atherosclerosis, we subset the main meta-analyzed reference to include only the pericyte-SMC-fibroblast level1 partitions. This subset was then reclustered using Seurat (v.4.1.0) with a resolution of 0.9 based on an additional silhouette width benchmark. Next, gene modules (encompassing top markers from differential expression analyses) specific to contractile (n=50), *Lgals3+* pioneer (n=50), and fibrochondrocyte (n=50) SMC phenotypes were extracted from a recent SMC lineage-traced murine scRNA meta-analysis. We also extracted a non-SMC-derived fibroblast module (n=50) as a negative enrichment control. Genes in each module were ranked by Log2FC and then converted to human homologs nomenclature and filtered to keep those with a one-to-one orthology relationship using custom wrapper functions with the biomaRt R package^108^ (v.2.46). We then calculated the enrichment of murine gene modules on individual cells within the pericyte-SMC-Fibroblast human subset using the UCell R package^64^ (v1.3.1). In addition to the enrichment of murine gene modules, we also obtained gene markers for each of the 17 SNN- derived clusters using the PrepSCTMarkers() and FindAllMarkers() functions from Seurat (v.4.1.0) setting the following thresholds: logFC=0.25 and min.pct=0.1. Final annotations for SMC subtypes were derived based on the UCell enrichment scores along UMAP coordinates and cluster markers.

### LD score regression applied to specifically expressed genes (LDSC-SEG) analyses

#### LDSC-SEG for SMC level 1 cell type annotations

Integration of scRNA and GWAS summary statistics was performed using the LDSC wrapper within the CELLECT python pipeline^103^. To improve the specificity of GWAS enrichment per cell type, we created a gene expression specificity matrix for level 1 annotations using the SCTransform-normalized expression matrix as input for the CELLEX python pipeline^103^. Shortly, gene expression specificity values (ESμ) output by CELLEX are derived using four different expression specificity metrics (Differential expression T-statistic, Gene enrichment score, Expression proportion and Normalized specificity index) and they represent a score that a gene is specifically expressed on a given cell type (level 1 annotation).

We downloaded GWAS summary statistics for: CAD (van der Harst et al)^35^; Myocardial infarction^36^; carotid intima-media thickness^37^, carotid artery plaques^37^, diastolic blood pressure, systolic blood pressure and pulse pressure from the Million Veterans Program ^38^; Alzheimer disease^39^; type 2 diabetes (UK Biobank)^40^; body mass index (UK Biobank)^40^; White blood cell count (UK Biobank)^40^. UK Biobank summary statistics were downloaded from https://alkesgroup.broadinstitute.org/UKBB/.

We used custom R scripts (https://github.com/MillerLab-CPHG/Human_athero_scRNA_meta) as well as the provided mtag_munge.py python script (https://github.com/pascaltimshel/ldsc/tree/d869cfd1e9fe1abc03b65c00b8a672bd530d0617) to convert GWAS summary statistics to a format compatible with that of the CELLECT S-LDSC wrapper. We then performed LDSC-SEG with the gene expression specificity matrix for level 1 annotations across the above described GWAS studies using the established CELLECT snakemake workflow as shown in https://github.com/perslab/CELLECT/wiki/CELLECT-LDSC-Tutorial.

#### LDSC for SMC level 2 cell type annotations

We proceeded to subset the whole meta-analyzed reference Seurat object to include only cells along the pericyte-SMC-Fibroblast partitions.

Metadata of this subset were used to generate the gene expression specificity matrix for level 2 annotations. In addition to GWAS studies described above, we also included summary statistics from our recent Coronary Artery Calcification (CAC) multi-ancestry GWAS meta-analysis^68^.

Munging of GWAS summary statistics and subsequent S-LDSC analyses were performed as described above.

### Cell communication analyses

Cell communication analyses were carried out using the Cellchat R package^69^ (v.1.5.0). We selected the CellChat human database (Interactions considered include secreted signaling, ECM-receptor and cell-cell contacts). First, we extracted SCTransform-normalized counts from the integrated Seurat object. For the first round of analyses, we separated cells from each disease status (lesion and non-lesion) and grouped them according to level 1 labels. We created a Cellchat object for matrices from each disease status using the createCellChat() function. We subsequently identified overexpressed genes in each condition using the identifyOverExpressedInteractions(). Communication probabilities were estimated with computeCommunProb() and aggregated cell communication networks calculated with the aggregateNet() function. We then merged lesion and non-lesion cellchat objects using the mergeCellChat() function. In order to identify pathways between Myeloid cells and SMCs that were enriched in each condition compared to the other, we input the merged Cellchat object to the function rankNet() with parameters (mode=”comparison, sources.use=”Macrophage”, targets.use=”SMC”). Significantly enriched pathways were denoted as those with *P*<0.05. To further explore differentially enriched pathways with increased granularity, we created a new CellChat object using normalized counts from Macrophages and SMCs from lesions and grouped them using their respective level 2 annotations. We computed communication probabilities and aggregated cell communication networks as described above. Circle plots for specific signaling pathways were generated with the netvisualAggregate() function. The top 30% of interactions (based on interaction weights/strength from computed communication probability) were used for plotting interactions between level 1-annotated cell types. Given that we had a larger number of cell types when deriving networks with level 2 labels, we chose to plot the top 15% of interactions.

### Pseudotime analyses for SMCs

Cells within the pericyte-SMC-fibroblast axis were subset to contain only contractile SMCs, transitional-ECM-SMCs, fibromyocytes and fibrochondrocytes. Single cell transcriptomic pseudotime analyses were performed using monocle3^70^ (v1.0.0). Given that gene expression within this subset was normalized, the SCTransform-normalized expression matrix and corresponding metadata were extracted from the corresponding seurat object. Metadata and SCT counts were used to create a cell_data_set object. In order to preserve clustering structure from previous analyses, we also extracted PCA/UMAP embeddings, cluster IDs and cell type annotations from the processed seurat object and inserted those into the corresponding slots of the cell_data_set object. A trajectory was then inferred using the learn_graph() and order_cells() functions setting contractile SMCs with the highest expression of *MYH11* as the root of the trajectory. DEG across the trajectory were calculated with grapth_test() and grouped into modules using the find_gene_modules() function. To model gene expression dynamics across pseudotime, we extracted pseudotime assignment values for each cell in the trajectory as well as SCTransform-normalized expression values and cell type annotations from the cell_data_set object. We then wrote a custom script to plot gene expression changes as a function of pseudotime where we applied cubic spline interpolation to expression values using the geom_smooth() function with parameters (method=“lm”, formula = y ∼ splines::ns(x, 3)).

### TF activity inference

For inference of TF activity, we also used a subset of the main reference only including SMCs, transitional SMCs, fibromyocytes and FCs. We downloaded a collection of curated TF regulons from the DoRothEA R package^75^ (v.1.8.0). We accessed human regulons using the dorothea_hs() function and only kept those with A, B and C confident scores for a more accurate prediction of regulon activity on each cell. Confidence scores had been previously defined based on the number of supporting evidence for each regulon^75^. TF activities for each cell were then estimated with the R package VIPER (v.1.24.0)^74^ providing the list of filtered regulons and the processed seurat object as input. Mean TF activities were then calculated across the SMC annotations of interest and the most variable TFs were selected for plotting.

### Human coronary artery tissue procurement

Freshly explanted hearts from orthopedic heart transplant recipients were obtained at Stanford University under approved Institutional Review Board (IRB) protocols with the respective informed consents. Hearts were arrested in cardioplegic solution and rapidly transported from the operating room to adjacent laboratory on ice. The proximal 5-6 cm of three major coronary arteries (LAD, LCX, RCA) were dissected from the epicardium, trimmed of surrounding adipose, rinsed in cold PBS and snap-frozen in liquid nitrogen. Human coronary artery tissue biospecimens were also obtained at Stanford University from non-diseased donor hearts rejected for orthotopic heart transplantation and processed following the same protocol as hearts for transplant. Reasons for rejected hearts included size incompatibility, risk for cardiotoxicity or comorbidities. Tissues were de-identified and clinical and histopathology information was used to classify ischemic, non-ischemic hearts and lesion and non-lesion containing arteries. All normal arteries originated from hearts with left ventricular ejection fraction (LVEF) greater than 50%. Frozen tissues were transferred to the University of Virginia through a material transfer agreement and IRB approved protocols.

### Coronary artery calcification GWAS meta-analysis data

The GWAS meta-analysis for coronary artery calcification (CAC) was conducted on 16 cohorts including 26,909 participants of European ancestry and 8,867 participants of African ancestry. CAC scores were calculated from computed tomography imaging at baseline, or first examination as described^68^. Genotyping quality control, imputation (1000 Genomes Phase 3), and variant filtering was performed as described. A joint meta-analysis of all available CAC GWAS was performed using a fixed-effects meta-analysis in METAL, using sample size weighted SNP p-values. The summary statistics from each study were combined using an inverse variance weighted meta-analysis.

### Pearson correlation calculations and gene set enrichment analyses

Normalized counts for cell types of interest were extracted from the corresponding Seurat object. Matrices were transposed to define genes as variables and then we calculated pairwise Pearson correlations for a gene of interest (e.g., *CRTAC1*) with all of the other genes across the cell types of interest using apply() and cor.test() functions with parameters (method=”pearson”) from the stats R package (v.4.0.3).

For gene set enrichment analyses, we calculated DE genes as described in the above section. We ranked genes by log2 fold change values (log2FC) and extracted the top 100 hits per cell annotation. We then use the gost() function within the R gProfiler2 package^105^ (v.0.2.1) with parameters (order=TRUE) to weight genes according to their log2FC values. We then selected significant GO:BP ontology terms (FDR < 0.05) and ranked them according to their adjusted *p*- values for plotting using custom functions from our scRNA_processing_utils.R script (https://github.com/MillerLab-CPHG/Human_athero_scRNA_meta). We found that the top GO:BP terms for fibrochondrocytes were highly redundant. Therefore, we used the gosemsim package^109^ (v2.16.1) and a custom script adapted from (https://github.com/YuLab-SMU/clusterProfiler/blob/master/R/simplify.R) in order to calculate semantic similarity between GO:BP terms. We removed highly redundant terms accordingly.

### Gene expression analysis in coronary artery datasets

#### RNA Extraction, QC, library construction and sequencing

Total RNA was extracted from frozen coronary artery segments using the Qiagen miRNeasy Mini RNA Extraction kit (catalog #217004). Approximately 50 mg of frozen tissue was pulverized using a mortar and pestle under liquid nitrogen. Tissue powder was then further homogenized in Qiazol lysis buffer using stainless steel beads in a Bullet Blender (Next Advance) homogenizer, followed by column-based purification. RNA concentration was determined using Qubit 3.0 and RNA quality was determined using Agilent 4200 TapeStation. Samples with RNA Integrity Number (RIN) greater than 5.5 and Illumina DV200 values greater than 75 were included for library construction. Total RNA libraries were constructed using the Illumina TruSeq Stranded Total RNA Gold kit (catalog #20020599) and barcoded using Illumina TruSeq RNA unique dual indexes (catalog # 20022371). After re-evaluating library quality using TapeStation, individually barcoded libraries were sent to Novogene for next generation sequencing. After passing additional QC, libraries were multiplexed and subjected to paired end 150 bp read sequencing on an Illumina NovaSeq S4 Flowcell to a median depth of 100 million total reads (>30 G) per library.

#### RNA-seq processing and analysis

The raw passed filter sequencing reads obtained from Novogene were demultiplexed using the bcl2fastq script. The quality of the reads was assessed using FASTQC and the adapter sequences were trimmed using trimgalore. Trimmed reads were aligned to the hg38 human reference genome using STAR^110^ (v.2.7.3a) according to the GATK Best Practices for RNA-seq. To increase mapping efficiency and sensitivity, novel splice junctions discovered in a first alignment pass with high stringency, were used as annotation in a second pass to permit lower stringency alignment and therefore increase sensitivity. PCR duplicates were marked using Picard and WASP was used to filter reads prone to mapping bias. Total read counts and Transcripts per million normalization (TPM) for both genes and isoforms was calculated from individual bam files using the RSEM (https://deweylab.github.io/RSEM/README.html) rsem-calculate-expression command with the paired-end option and gencode version 32 as a reference^111^.

### Coronary artery proteomics data generation and analysis

#### Tissue processing

Frozen human coronary artery segments were shipped in 1.5 mL microcentrifuge tubes to King’s College London (London, United Kingdom). To remove glycans attached to ECM proteins, we used deglycanation enzymes (Heparinase II (Sigma-Aldrich H6512-10UN), Chondroitinase ABC (Sigma-Alrich C3667-5UN), Keratanase (G6920-5UN)) and a glycoprotein deglycosylation kit (Merck catalog #362280). We then used Water-18O (97% atom) to label N-linked glycosylation sites. After deglycosylation the ECM protein samples (n=150) underwent denaturing, reduction, alkylation, precipitation, and overnight trypsin digestion. We purified the resultant ECM fragments with AssayMAP C18 cartridges (Agilent) on an Agilent Bravo AssayMAP robot. We analyzed the purified peptide samples using nanoflow liquid chromatography tandem mass spectrometry (LC-MS/MS). We performed data-dependent analysis (DDA) (on the top 15 ions in each full MS scan) using a nanoflow LC system (Dionex UltiMate 3000 RSLC nano) coupled to a high-resolution accurate-mass Orbitrap mass analyzer (Q Exactive HF, Thermo Fisher Scientific).

#### Proteomics data analysis

We used the Thermo Scientific Proteome Discoverer software (v.2.3) to search the raw proteomic data files against the human database (UniProtKB/Swiss-Prot version 2019_01, containing 20,349 protein entries) using the Mascot server (version 2.6.0, Matrix Science). We measured protein abundance in each sample using label-free quantitation (LFQ). Data was analyzed according to the King’s College London pipeline and processing protocol^112, 113^. Data was normalized according to the total ion intensity and subsequently scaled to remove batch effects. We filtered out proteins with more than 30% missing values. For the remaining missing values, we performed imputation with the K-nearest neighbor (KNN) impute algorithm. To tune the parameter k of the KNN-impute method we experimentally tested the Euclidean distance of the imputed values compared to the real ones for 100 randomly selected values, testing for k=2 until 20. The optimal k value was set to 5 according to this procedure and this was applied to impute all the remaining missing values. Values were then displayed in Log2 scale.

#### Definition of disease categories for transcriptomic and proteomics comparisons

Disease status of coronary artery segments was determined based on our previous publication^76^. For the present study, samples that are lesion-free (no evidence of atherosclerosis) or harbored subclinical/early lesions were included within the “non-lesion group”. Samples with evidence of advanced/complex atherosclerotic lesions were included in the “lesion” group for comparative transcriptomics and proteomics analyses.

### Tissue metadata for bulk transcriptomics and proteomics analyses

Mean age for individuals in transcriptomics comparisons = 52.1 years. From these individuals, 14 (29%) were females and 34 (71%) males. Mean age for individuals in proteomics comparisons = 51.2 years. From these individuals, 16 (29%) were females and 40 (71%) males.

### STARNET regulatory networks and clinical trait enrichment analysis

Based on STARNET^86^ multi-tissue bulk RNA-seq data, tissue specific and cross-tissue co-expression modules were inferred using WGCNA^114^. Enrichment for clinical traits was computed by aggregating Pearson’s correlation *P* values by co-expression module using Fisher’s method. Enrichment for DE genes was calculated with the hypergeometric test using DESeq2 called genes (30% change, FDR <0.01) adjusting for age and sex covariates. The gene regulatory network for *CRTAC1* and *LTBP1* co-expressed genes was inferred using GENIE3^115^. Weighted key driver analysis was then applied to identify hub or highly influential genes in the regulatory network using the Mergeomics R package^116^.

### Histological analysis of human coronary artery tissues

Human coronary artery tissues were obtained as described in the above section. Briefly, coronary artery segments were isolated from control/subclinical and advanced atherosclerotic left and right coronary artery branches. Tissues were embedded in OCT blocks, snap-frozen in liquid nitrogen and stored at −80 °C. Segments for histological staining were chosen based on sample availability as well as the degree of correlation between RNA/protein levels. OCT embedded human coronary artery segments were cryosectioned at −20 °C and 8-μm thickness. A minimum of two sections per sample were placed on each slide and then blindly stained with: Oil Red O (ORO), Picro-Sirius red (PSR) and Hematoxylin and Eosin (H&E) at the UVA Research Histology Core. Briefly, for ORO staining, frozen sections were fixed in 10% Neutral Buffered Formalin solution, washed, and stained in Oil Red O solution (Poly Science #s2120) for 5 min. After washing, slides were stained in Hematoxylin solution (Richard Allen #7221) for 1 min before rinsing and mounting with aqueous mounting medium. For H&E, slides were stained using Hematoxylin 360 reagents manufactured by Leica in an automated Gemini Stainer. For PSR staining, slides were placed in Picro-Sirius red solution (Direct Red 80 in saturated aqueous solution of picric acid) for 1 hour, rinsed in deionized water and washed twice in acidified water. Slides were then dehydrated in ethanol, cleared in xylene, and mounted. Whole slide images were then captured at approximately 100,000 x 30,000-pixel resolution using a Hamamatsu NanoZoomer S360 Digital Slide Scanner C13220 at the Biorepository and Tissue Research Facility at UVA.

### Immunofluorescence of human coronary artery tissues

We performed immunofluorescence staining in tissue sections adjacent to those used for histology. Sections were processed for immunostaining as follows: Sections were retrieved from the −80 °C freezer and allowed to briefly come to room temperature for 1 min. Sections were then rehydrated in 1X PBS and then fixed in 4% formaldehyde for 10 min at room temperature. This was followed by three PBS washes. Sections were then permeabilized using 0.1% Triton-X-100 for 10 min at room temperature. Upon permeabilization, sections were washed with PBS and then protein blocking was done with 10% normal donkey serum for 1 h at room temperature. After blocking, slides were incubated with an anti-LTBP1 rabbit polyclonal antibody (Proteintech, 26855-1-AP; 1:250 in antibody dilution solution (1% BSA)) overnight at 4 °C or no antibody negative control (antibody dilution solution). Optimal dilution concentration was determined with previous titrations with control tissues. Each slide had at least two sections stained with primary antibody and one section used for the negative control. Sections were washed with PBS and incubated with donkey anti-rabbit Alexa Fluor 555 conjugated secondary antibody (Thermo Fisher, A31572; 1:1000) and aSMA conjugated to FITC (Sigma, F3777; clone 1A4; 1:500) for 1 h at room temperature. Slides were then washed with PBS and nuclear counter-staining was performed with DAPI (0.1 ug/ml) (MiltenyiBiotec, 130-111-570). Slides were subsequently coverslipped with aqueous mounting media. Stained sections were visualized using an Olympus BX41 microscope under x10 and x20 objective magnifications.

Images were obtained using a Nikon color camera DS-Fi3 at an NIS Elements imaging software (v110.04.3707.E9).

### Coronary artery snATAC-seq tissue processing and data analysis

#### Coronary artery sample processing and nuclei isolation for snATAC

snATAC-seq data for human coronary arteries was generated as described in our previous publication^76^. Briefly, isolated nuclei were processed using the 10X Genomics Chromium Single Cell ATAC and fastq files were preprocessed using the 10X Genomics Cellranger pipeline (CellRanger ATAC v1.2.0) using the hg38 reference genome and default parameters.

#### Dimensionality reduction, clustering of snATAC-seq data and generation of gene activity scores

Outputs from Cellranger were analyzed with the ArchR pipeline^77^ (v.1.0.2). Fragment files for each of the 41 patients were used to generate ArchR arrow files. We filtered low-quality cells with the following parameters: TSS enrichment > 7, unique number of fragments > 10000 and a doublet ratio < 1.5. The genome was then divided into 500bp windows and then fragments within each window were used to generate a tile matrix (28316 cells x ∼ 6 million tiles). Iterative latent semantic index (LSI) was then used to reduce dimensionality of the tile matrix. We checked for batch effects using Harmony (v.1.0) and did not observe major differences in the data clustering structure (clusters driven by individual samples). We then used the first 30 components output by LSI for running non-linear dimensionality reduction (UMAP). Subsequent cell clustering was performed using the SNN modularity optimization-based algorithm as implemented in Seurat (v.4.1.0). Chromatin accessibility (defined as the number of fragments within each tile) within gene bodies as well as proximally/distally from the TSS was used to infer gene expression by means of a gene activity score model. In this model, the number of fragments inside tiles of gene bodies are considered as well as surrounding tiles. To account for the activity of putative distal regulatory elements, an exponential weighting function is applied where tiles that reside further from genes TSS are assigned lower weights. Additionally, this model imposes gene boundaries to minimize the contribution of unrelated regulatory elements to a specific gene score.

#### TF motif enrichments

Enriched TF motifs for each cell type were predicted using the addMotifAnnotations() function in ArchR. Z deviation scores for each TF were then estimated with the chromVAR R package^117^ (v.1.12.0).

## Statistical methods

Differentially expressed genes for the scRNA-seq data were calculated using a Wilcoxon Rank-Sum Test as implemented in Seurat (v.4.1.0). Statistics for bulk transcriptomics and proteomics data were performed using the stats R package. To compare levels of normalized gene expression in human coronary arteries across disease conditions (non-lesion vs lesion), we used a non-parametric Wilcoxon Rank Sum Test. To analyze protein levels across the above-mentioned disease conditions, we used an Student’s *t*-test (two-sided). Normality of the data was checked by generating quantile-quantile plots. Variance of the data was inspected by plotting the distribution of gene expression and protein values across samples using a density plot. Wilcoxon Rank Sum Tests and Student *t* tests were performed with a significance threshold of *p* < 0.05. The number of samples used in each analysis is indicated within each figure legend. The Pearson’s product moment correlation coefficients for comparing normalized gene expression and protein levels were calculated using cor.test() within the stats R package. For calculations of Pearson correlations across the entire transcriptome in the scRNA-seq data, *p*-values from were adjusted for multiple testing using the Benjamini Hochberg correction as implemented in the stats package with the p.adjust() function and parameters (method=”fdr”).

## Data Availability

Raw count matrices included in this study were accessed through GEO and Zenodo. Raw count matrices for *Wirka* et al.^18^, *Pan* et al.^19^, *Alsaigh* et al.^16^ were obtained through the following accession numbers: *Wirka et al* (GSM3819856, GSM3819857, GSM3819858, GSM3819859, GSM3819860, GSM3819861, GSM3819862, GSM3819863); *Alsaigh et al* (GSM4837523, GSM4837524, GSM4837525, GSM4837526, GSM4837527, GSM4837528); *Pan et al* (GSM4705589, GSM4705590, GSM4705591). Raw count matrices from *Hu* et al.^17^ were obtained from Zenodo (https://zenodo.org/record/6032099#.Y1RDa-zMITU). The corresponding accession numbers can also be found in **Supplementary Table 1**. Bulk RNA-seq data from human carotid lesions^85^ was accessed through GEO with the accession number GSE120521.

The integrated reference will be placed in plaqview for public visualization. The Seurat object with annotated clusters will be made available upon request.

## Code Availability

Code used for processing of raw count matrices, integration benchmark and other downstream analyses can be found in the following Github repository: https://github.com/MillerLab-CPHG/Human_athero_scRNA_meta

## Acknowledgements

This work was supported by grants from: the National Institutes of Health (grant numbers R01HL148239 and R01HL164577 to C.L.M.; R35GM133712 to C.Z.; R01HL142809 and R01HL159514 to R.M.; K01HL164687 to C.L.L.C.; R01HL125863 to J.L.M.B; R01HL130423, R01HL135093 and R01HL148167-01A1 to J.C.K.), the American Heart Association (grant number 20POST35120545 to A.W.T.; AHA909150 to J.V.M.; A14SFRN20840000 to J.L.M.B.), the Swedish Research Council and Heart Lung Foundation (grant number 2018-02529 and 20170265 to J.L.M.B.), the Fondation Leducq (grant number ‘PlaqOmics’ 18CVD02 to C.L.M., S.W.vdL., J.L.M.B., and M.M.), the European Union H2020 TO_AITION grant number 848146 (to SWvdL) and the Single-Cell Data Insights award from the Chan Zuckerberg Initiative, LLC and Silicon Valley Community Foundation (to C.L.M., C.Z. and S.W.vdL.). We would like to thank Dr. Timothy Bullock and Dr. Sanja Arandjelovic for their tremendously helpful insights regarding myeloid and lymphoid cell subtype annotations.

## Author contributions

C.L.M. supervised research primarily related to the study. M.M., J.C.K., J.L.M.B., R.M., N.C.S., C.Z., and S.W.vdL. jointly supervised research secondarily related to the study. J.V.M. and C.L.M. conceived and designed the experiments. J.V.M., G.A., A.W.T., K.T., and C.L.L.C. performed the experiments. J.V.M. performed the statistical analyses. J.V.M., A.W.T., C.J.H., and K.T. analyzed the data. M.B., M.K., P.P., M.M., J.C.K., J.L.M.B. and S.W.vdL. contributed reagents/materials/analysis tools. J.V.M., D.W., G.A., A.W.T., C.J.H., N.C.S. and C.L.M. wrote the paper.

## Competing Interests

J.L.M.B. is a shareholder in Clinical Gene Network AB who have a vested interest in STARNET. S.W.vdL. has received funding from Roche for unrelated work. C.L.M. has received funding from AstraZeneca for an unrelated project. R.M. has received funding from Angea Biotherapeutics and Amgen and serves as a consultant for Myokardia/BMS, Renovacor, Epizon Pharma, and Third Pole, all unrelated to the current project. J.C.K. is the recipient of an Agilent Thought Leader Award, which includes funding for research that is unrelated to the current project. The other authors declare no competing interests.

## Notes

### Summary of Updates

We provide updated text and data supporting our main findings on a unified single-cell map of human atherosclerosis.

